# Moving Beyond the Mean: Subgroups and dimensions of brain activity and cognitive performance across domains

**DOI:** 10.1101/2020.02.28.970673

**Authors:** Colin Hawco, Erin W. Dickie, Grace Jacobs, Zafiris J. Daskalakis, Aristotle N. Voineskos

## Abstract

Human neuroimaging during cognitive tasks has provided unique and important insights into the neurobiology of cognition. However, the vast majority of research relies upon group aggregate or average statistical maps of activity, which do not fully capture the rich variability which exists across individuals. To better characterize individual variability, hierarchical clustering was performed separately on six fMRI tasks in 822 participants from the Human Connectome Project. Across all tasks, clusters ranged from a predominantly ‘deactivating’ pattern towards a more ‘activating’ pattern of brain activity, with differences in out-of-scanner cognitive test scores between clusters. Cluster stability was assessed via bootstrapping approach. Cluster probability did not indicate distinct/clear clustering. However, when participants were plotted in a dimensionally reduced ‘similarity space’ derived from bootstrapping, variability in brain activity among participants was best represented multidimensionally. A ‘positive to negative’ axis of activity was the strongest driver of individual differences.

## Introduction

Functional magnetic resonance imaging (fMRI) has resulted in a dramatic improvement in our understanding of human brain function and cognition over the past 25 years. The great majority of these studies have made conclusions about groups of individuals (based on a disorder, age-range, or lack of a disorder, i.e. a healthy group). These findings can give the impression that any ‘group’ is homogeneous, i.e. brain activation representing a group-average might represent that of an individual. However, a growing body of work is bringing to light the tremendous variability among individuals in brain function. For example, typical patterns of connectivity observed at the group level are not well represented at the individual level^1^, and individual variability is greater in heteromodal cortices such as the fronto-parietal network^2^. The extent and distribution of different patterns of activity among individuals during cognitive tasks is not optimally captured by group statistical maps^3-5^. Further, there is evidence to support that people may use different cognitive strategies during task performance^6-8^. For example, strategy training in individuals with normal memory abilities can develop patterns of activity that resemble exceptional ‘memory athletes’^9^. Additionally, individuals with greater differences in cognitive styles have concomitant differences in brain activity^6^, demonstrating a relationship between cognitive strategies during task performance and patterns of brain activity.

Much progress has also been made in the identification of individual brain systems, which vary according to dimensions such as cognitive abilities. Regression approaches often used in these investigations are predominantly sensitive to variation, which linearly maps onto a common spatial pattern of brain-behavior relationships across all individuals. However, such approaches are not designed to capture unique brain-behavior relationships among sub-groups of participants^10^. There are also important statistical considerations whereby linear regressions may overestimate brain-behavior relationships^11^, and issues regarding reliability at lower sample sizes have been well-documented^12,13^. Due to the challenges of interpreting single subject data related to the high dimensionality and complexity of brain imaging data, few fMRI studies have focused on mapping differences in brain function among subgroups, and even fewer among individuals^4,6,14,15^. One intermediate approach between group level analyses and visualizing data at the individual level is to perform clustering on participants, grouping together those with similar neurobiological characteristics^3,16-19^. In a recent investigation including individuals with and without psychiatric disorders, we identified and replicated three sub-groups (clusters) demonstrating distinctly different patterns of brain activity during emotional face processing. These sub-groups were comprised of a ‘deactivating’ group, a typical/expected group (i.e. representative of the group average), and a ‘hyperactivating’ group^3^. These sub-groups were not related to diagnostic status (e.g. with or without psychiatric diagnosis), but did show differences in cognitive performance. It remains unknown whether a) such patterns exist across most fMRI tasks, and b) if such variability would be optimally represented as discrete sub-groups or along a continuum of individuals.

The purpose of this study was to better identify the different patterns of human brain activity during task processing across a range of cognitive domains. We were also interested to determine whether these patterns could be best represented by discrete subgroups, or whether they exist along dimensions at an individual level. Finally, we were interested to determine whether these new patterns of brain function among subgroups and individuals would relate to cognitive performance. We leveraged the large sample available in the Human Connectome Project (HCP), which allowed us to examine our question using all fMRI tasks covering six cognitive domains^20^, namely Emotional, Gambling, Language, Relational, Social, and Working Memory. Furthermore, cognitive performance data from those tasks were also available allowing us to consider how different patterns in brain function were related to performance.

## Results

### Clustering revealed a gradient of activity across individuals for each task

Data from 822 participants from the HCP900 release who had completed all six fMRI tasks were included^20^. For each participant for each task, data were extracted from the t-maps from the minimal processing pipeline with 8mm surface smoothing, provided by the HCP. For each task, hierarchical clustering with Euclidean distance and Ward’s linkage^3,21^ was applied for k=2 through k=10, with k indicating the number of clusters in a given solution. Subsequently, a group analysis (one sample t-test) was run in SPM12 to examine group patterns of activity within each cluster for each task. The aim of this study was to visualize the range of activity across sub-groups of participants rather than identify specific ‘best’ cluster solutions. For the initial cluster solution, results from k=4 are presented in Figure 1, as a representative visualization of the result. Results from all other values of k are presented in Supplemental Figures 1-8, as well as being available online (see code and data availability in methods). A similar pattern was observed at other values of k (i.e. a range of activity from most deactivating to most activating). Participants separated along a gradient from predominantly negative activation (or deactivation) towards predominantly positive activation. While the center/intermediate clusters tended to partially resemble the full sample group average, the more ‘deactivating’ and ‘hyper-activating’ clusters (shown on the top and bottom in Figure 1, respectively) showed distinct patterns from each other and the total sample group maps. Exemplifying this were groups with strong positive and strong negative activation in the Gambling and Relational tasks. These findings demonstrate substantial variations in patterns of brain activity across groups, which were not captured by a standard analysis using the full sample. In order to confirm that these group patterns ranging from deactivation to activation were not specific to hierarchical clustering, clustering was repeated using the K-means approach (k=4), which produced similar patterns of activation across clusters (Supplemental Figure 9).

**Figure 1:**
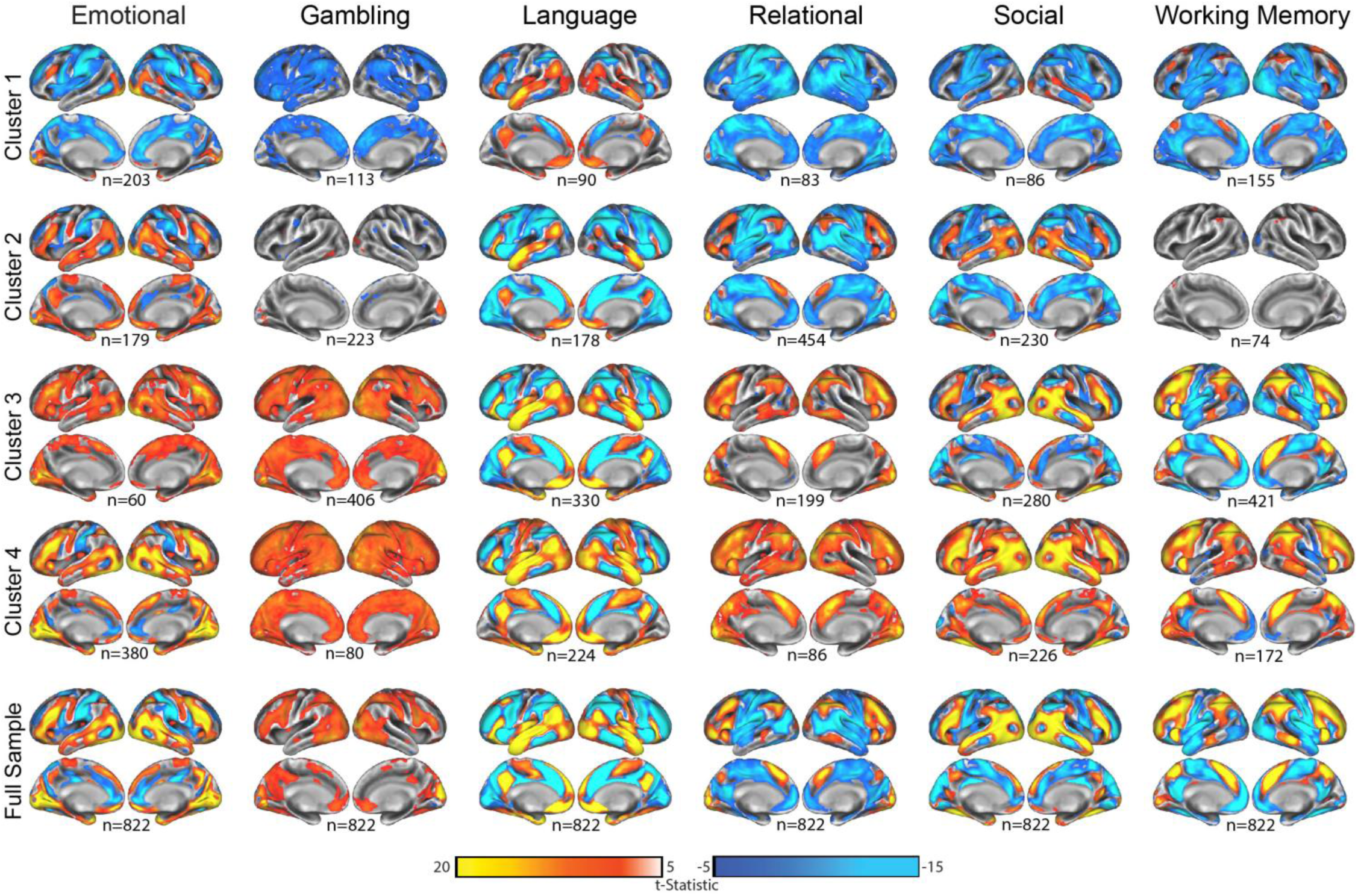
Group analyses for each task (columns) by cluster (top four rows) for k=4, and for comparison across the whole sample (bottom row). For visualization, clusters are arranged from most negatively activating (top) to most positively activating (bottom). All maps are thresholded at t ≥ 5 for visual comparisons. Group analyses (one-sample t-tests) were completed in SPM12. The number of participants in each cluster is presented below each map. The left hemisphere is shown on the left for each image.

### Clustering of participants is not similar across tasks

Previous work has suggested task activity is built on an architecture of functional connectivity^22-24^, indicating that participants with similar functional connectivity may show similar task activation patterns. In order to assess if participants clustered similarly across the six tasks, we used the Adjusted Rand Index (ARI). The ARI indicates similarity between two cluster solutions while adjusting for random chance (i.e. ARI = 1 indicates perfect overlap in cluster membership, while ARI ∼ 0 indicates no relationship, random chance). ARI was calculated for each pair of tasks (e.g. Emotional-Gambling, Emotional-Language, etc.) for each value of k. Across all pairs of tasks at all levels of k, the ARI was very close to zero (all ARI < 0.03), demonstrating there was no relationship between how participants clustered across tasks.

### Clusters show differences in cognitive abilities

The behavioral relevance of clusters was considered by examining behavioral/cognitive differences among clusters using scores from cognitive tests within the HCP battery (Figure 2). For each task and each clustering solution from k=2 to k=10, a one-way ANOVA was performed for each of 12 Z-transformed test scores to examine cognitive differences between clusters. The p-values were FDR corrected (648 total tests; FDR corrected threshold was p < 0.0188) across all ANOVAs performed (Figure 2). All tasks showed at least some significant differences in cognitive scores among clusters, with the Language, Relational and Working Memory tasks showing the strongest relationships, while clusters in the Gambling task only differed in picture sequence and flanker cognitive scores. In some cases, the p-values for cluster differences were quite small (e.g. p < 1×10^−10^), though effect sizes were small (largest effect size, eta^2^ = 0.112). As a summary visualization, cluster differences in performance on Raven’s progressive matrices (PMAT; proposed to approximate a test for overall fluid intelligence^25^) for task showing significant cluster differences in PMAT are plotted in Supplemental Figure 10. While in our previous work the ‘deactivating’ cluster showed the strongest cognitive capabilities^3^, here it was observed that the pattern of activity associated with the highest PMAT score varied by task. A similar analysis was done using behavioral scores (e.g. accuracy or reaction time) taken from performance on the fMRI tasks, representing performance more closely related to the tasks themselves (Supplemental Figure 11). For all tasks except Gambling, group differences were found between clusters in behavioral performance on the fMRI task.

**Figure 2:**
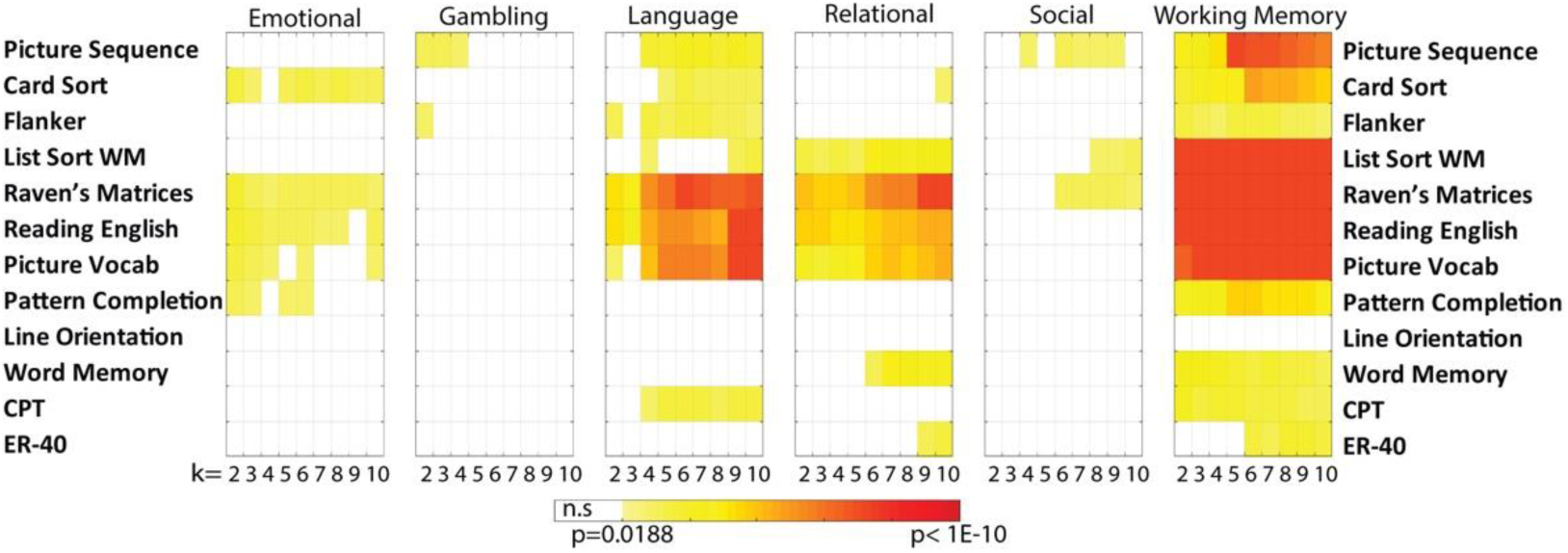
Results of one-way ANOVAs comparing differences between clusters on cognitive test scores. For each fMRI task a one-way ANOVA was run for each cluster solution (k=2 to k=10; columns) for each cognitive test (rows; test names are shown on both sides for readability). P-values from each ANOVA are presented as a colored box. ANOVAS were run for the clustering solutions using hierarchical clustering. Any p-values which were not significant (FDR corrected across all tests, p<0.05) are shown as white.

### Bootstrapping stability assessments suggest our apparently distinct sub-groups are ‘pseudo-groups’, which fall along a continuum

A key goal of the present study was to better understand if participants fall more along a continuum or into relatively discrete groups (i.e. clusters). In order to interrogate the underlying structure of the cluster solutions, a bootstrap without replacement approach was employed. For each task and cluster solution from k=2 to k=10, 1000 bootstraps were run using a randomly selected 75% of participants to recalculate clusters. For each combination of task and k value (e.g. Working Memory task, k=4), the proportion of times that any given pair of participants clustered together whenever they were both present in a given bootstrap solution was calculated (i.e. showing the probability of any two participants clustering together across bootstraps). This was then organized as an NxN (822×822) ‘cluster probability’ matrix (Figure 3). In the case of highly stable clustering, we would expect a “box-like” pattern within these matrices; if the range of patterns of brain activity were best represented by a pure spectrum, we would expect a wide diagonal blob shape with no bulges. Instead, the patterns emerged between these two outcomes; a spectrum across the middle with varying degrees of off-diagonal bulges. This suggests that participants do not fall into distinct sub-classes, or the borders between classes are not easily distinguishable, while the off-diagonal bulges suggest some localized clustering representing sub-groups along the continuum, to varying degrees across tasks.

**Figure 3:**
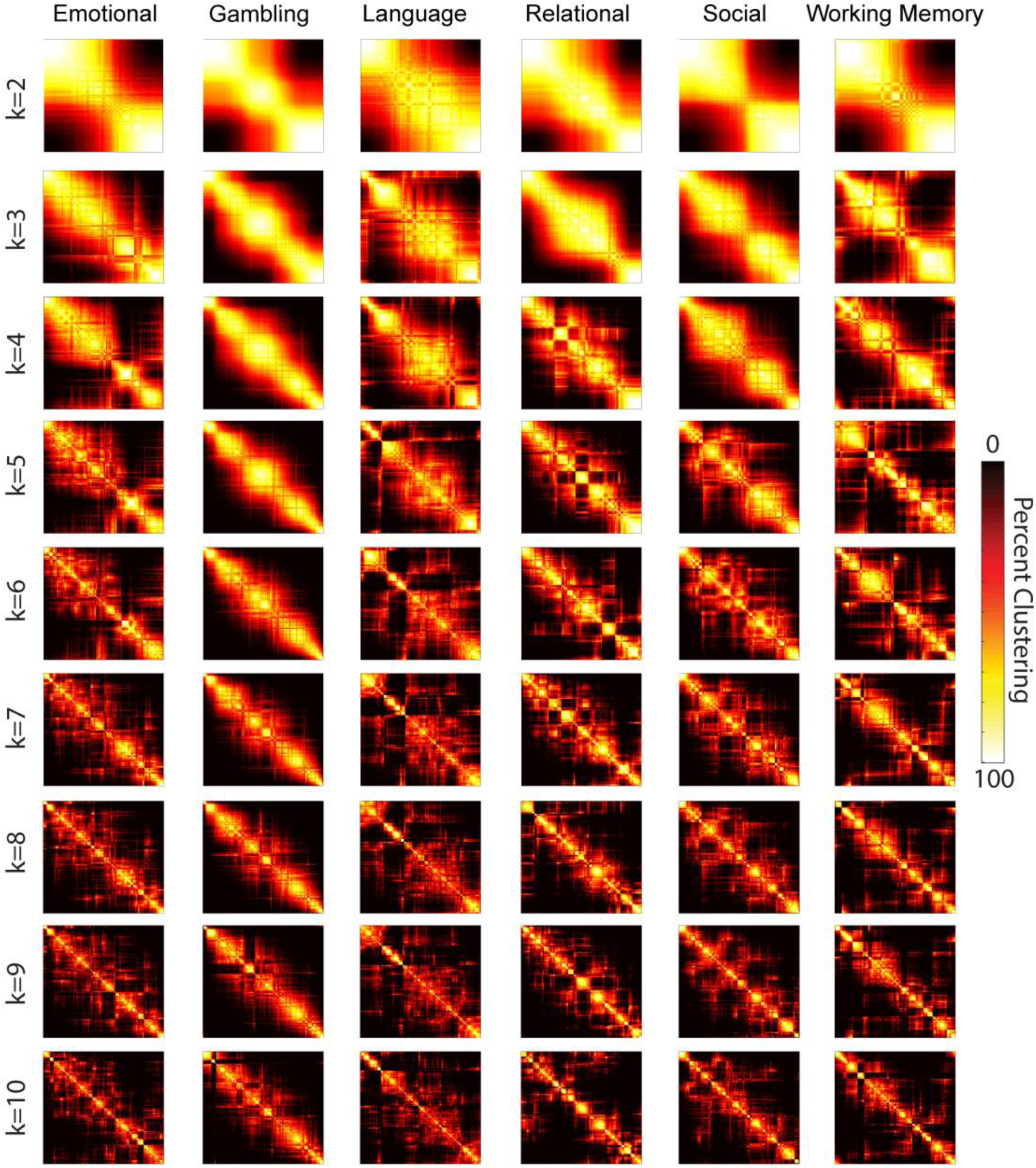
Cluster probability matrices for each task modality, k=2 to k=10. The matrices are each an NxN (N being participants, 822 total), showing the percentage of times that any pair of participants clustered together if both were present in the bootstrap. Each matrix was individually reordered to maximize the sum of the similarity between adjacent participants (MATLAB’s optimal leaf order function). If participants formed discrete clusters we would expect to see ‘squares’ within the matrix.

### Component scores of the clustering probability matrices reveal a new manifold space of participant similarity

The cluster probability matrices (Figure 3) suggest some discrete clustering in cases (e.g. off diagonal bulges in the graph), but there was an apparent dimensional component in all cluster solutions. In order to further explore these results, a principal component analysis (PCA) was performed on each cluster probability matrix to identify underlying patterns observed across tasks, such as in Figure 4A. The result was plotted and examined for all solutions. The k=4 data were further analyzed as a representation of all patterns because it maximized the shared variance across the three components (i.e. the top three components accounted for 90% or more of the variance in each task; Figure 4B).

**Figure 4:**
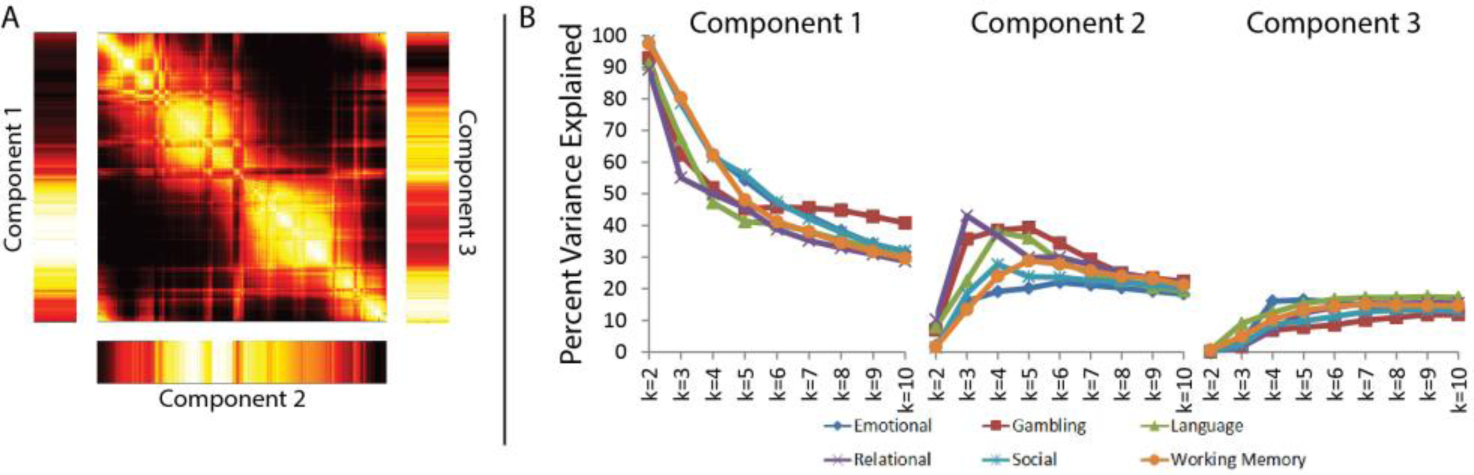
A) Example components from the PCA of the bootstrap permutation matrix for the Working Memory task (k=4). The cluster probability matrix is shown in the center while the bars represent the top three components of the PCA (brighter colors indicate that the participant scores higher in that component). B) Percentage of variance explained by the components 1, 2, and 3 for the PCA of the cluster bootstrap matrices for all six tasks, for k=2-10.

Component scores were visualized by plotting the location of each participant on a 3D scatterplot based on participants’ scores from the first three components (Figure 5, left panel; results for all values of k are shown in Supplemental Figure 12). Two specific shapes emerged: either a ‘snake’ shape, with two curves (for Gambling, Relational, and Social), or a folded circular shape (a ‘tortilla chip’ shape; for Emotional, Language, and Working Memory), in which all participants fall on or near the surface of the ‘tortilla’ shape. Further, at higher values of k, the scatterplots for the Relational and Social tasks transition from a ‘snake’ to a ‘tortilla’ shape (Supplemental Figure 12), suggesting these two shapes may share a common underlying derivation. While the plots show the multi-dimensional complexity underlying similarity across participants that may drive clustering, the ‘snake’ shape suggests that some tasks may fall along a lower-dimensional continuum. In order to further explore this, component scores were plotted color-coded by mean activity during the task, defined as the mean of all t-statistics within the cortex for each participant for the contrast map used in the clustering analysis. Position within both ‘snake’ and ‘tortilla’ shapes were related to mean activity; in ‘snake’ shapes, mean activity related to order along the ‘snake’, while for ‘tortilla’ shapes, participants varied in mean activity in a continuum along the circular edge (Supplemental Figure 13). The space captured within these 3D plots may represent a manifold of similarity in patterns of brain activity, which underlie individual variability across the sample.

**Figure 5:**
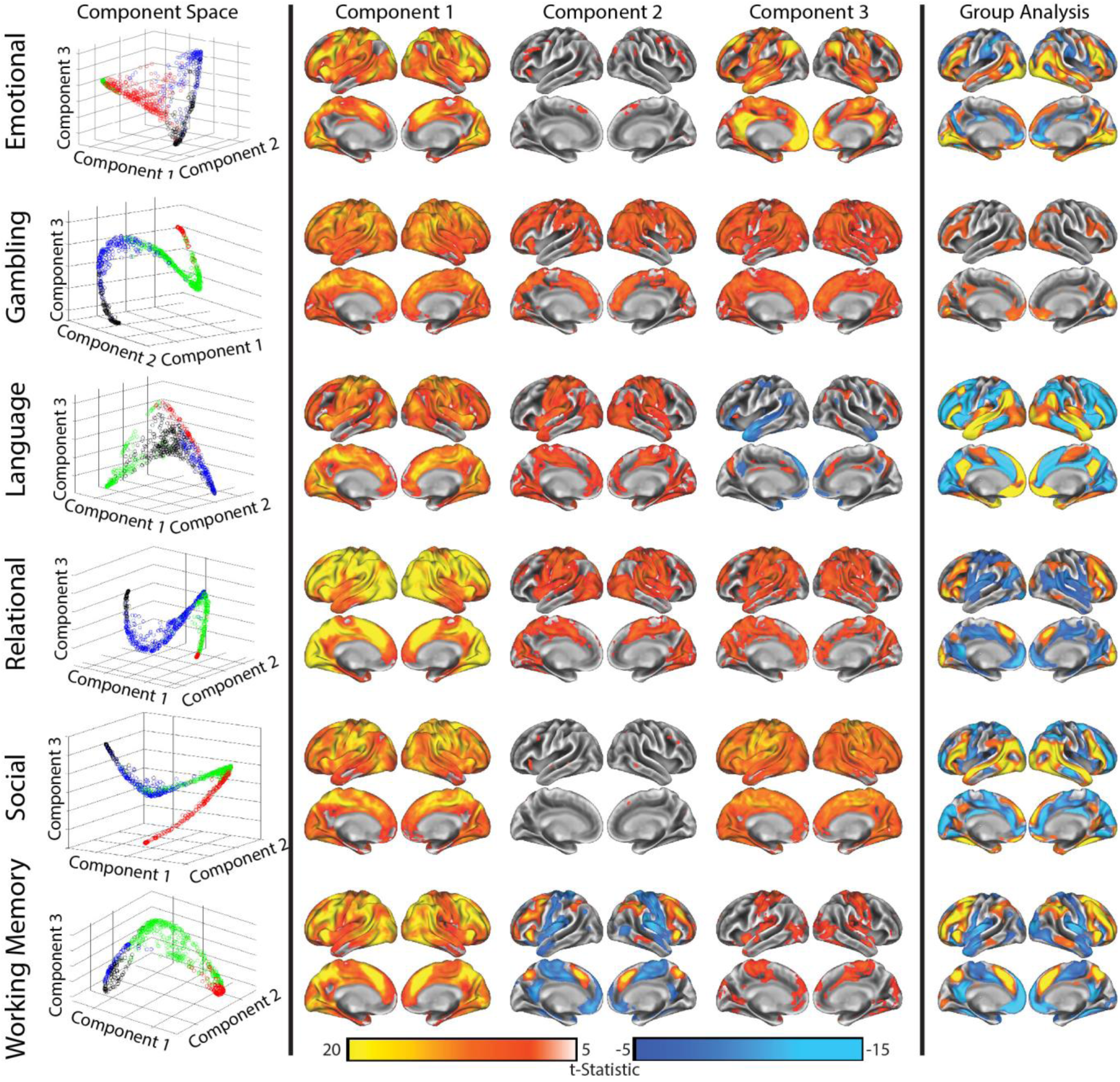
Left panel: participants mapped into 3D space based on their scores in the first three components of the cluster probability matrix. Each point represents one participant. Colors show the full sample cluster solution (as in Figure 1), with cluster 1 (most deactivating) as black, cluster 2 as blue, cluster 3 as green, and cluster 4 (most positive activating) in red. Center panel: the results of a regression analysis of component scores on brain activity, showing regions in which participants with a higher or lower brain score had greater activity. Right panel: group statistical analysis by task using the whole sample, as in Figure 1, shown for visual comparison with regression results.

In order to consider whether the use of component scores was an appropriate means of data reduction of the bootstrap probability matrix, multi-dimensional scaling was applied to the cluster probability matrix to calculate 3 dimensions; multiscale dimensions correlated with component scores from 0.89 to 0.99, and the ‘snake’ or ‘tortilla’ shapes were recreated (Supplemental Figure 14), suggesting the ‘snake’ or ‘tortilla’ shapes were not merely a function of using PCA.

### Patterns of brain activity linked to clustering probability components

The patterns observed in the cluster probability matrices and subsequent mapping onto clustering principal component space are driven by variations in the spatial brain activity across individuals. In order to visualize the spatial patterns related to the components, we used a regression approach to identify regions which had more or less activity in participants with a higher component score. Results of the group whole brain regression analysis using k=4 for each task are shown in Figure 5 (center panel). For all six task modalities, the first component mapped onto global activity with the strongest relationships in regions showing positive activity in the whole sample group maps (whole sample group maps are shown in Figure 5, right panel). Components 2 and 3 varied by task and were driven by specific variations in activity in task relevant networks.

In order to further quantify the regions showing the strongest relationships with component 1 (i.e. determine if specific brain systems had stronger loading onto this component), we extracted the mean t-value for six networks from the Yeo 7-network parcellation^26^ (default mode, fronto-parietal, dorsal attention, salience/ventral attention, visual, and sensory-motor networks; the ‘limbic’ network was excluded due to poor signal quality in those regions) for the whole brain regression analysis of principal component 1 of the cluster probability matrix (as shown in Figure 6). The dorsal attention network showed the strongest relationship with component 1 for the Emotional, Gambling, Relational, and Social tasks. In contrast, the salience network in Language and fronto-parietal network in Working Memory had the highest mean t-values, although the dorsal attention network was still prominent. The default mode network had the lowest t-values across all six tasks. This shows that while component 1 reflects a pattern of global brain activity, it is nevertheless weighted towards the ‘task-positive’ networks^27,28^.

**Figure 6:**
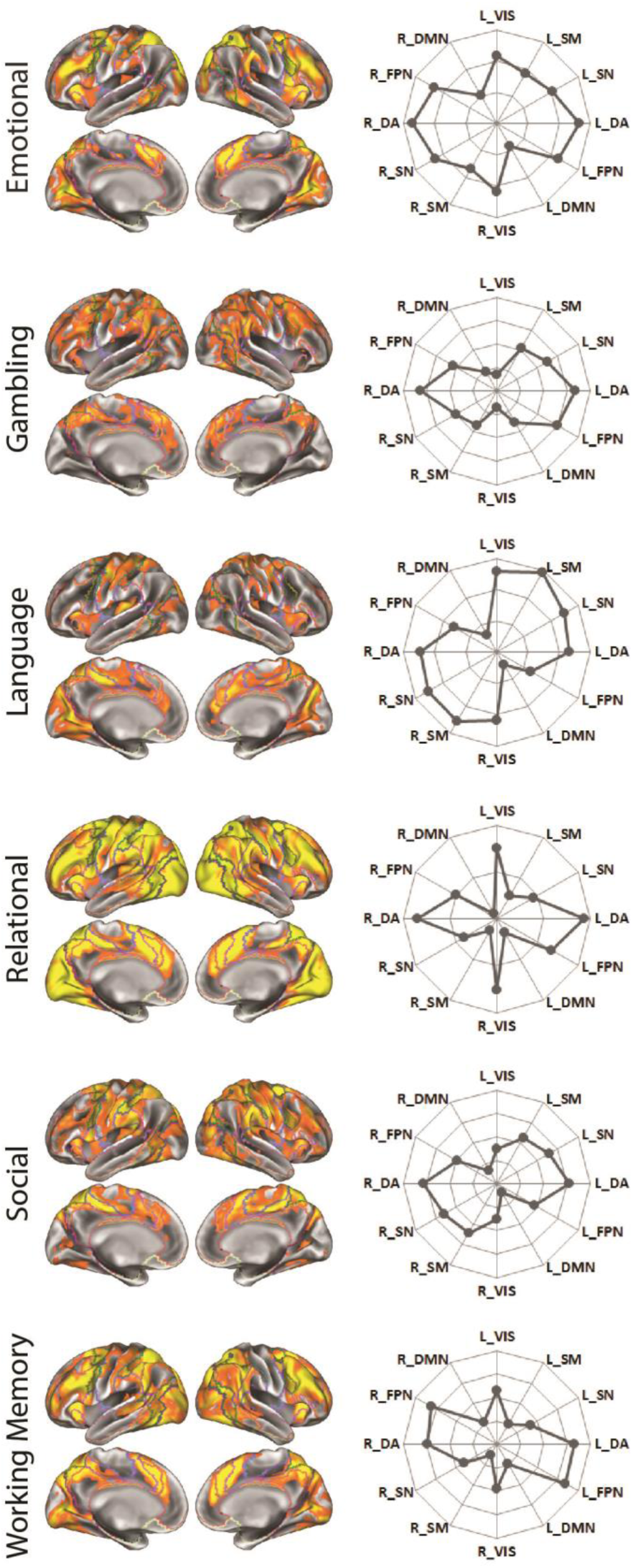
Network loadings of the regression analysis of component 1 on task activity. The left panel shows the regression t-statistics for each task thresholded at t>=10 to better show regions with the most significant relationships. Outlines on the brain surfaces represent the six networks from the Yeo 7 network parcellation^26^. The radar plots (right) show the relative contribution of each of the six networks, expressed as the average t-statistics within all vertices across that network (higher values on the outside). Mean t-stats were demeaned across networks to emphasize relative differences (Vis=Visual, SM=sensory-motor, SN=salience/ventral attention, DA=dorsal attention, FPN=fronto-parietal network, DMN=default mode network, L=left, R=right).

### Cognition is linked to clustering probability components

We performed Spearman’s correlations between the three components and out-of-scanner cognitive scores (Figure 7). No significant correlations were observed in the Social task, but were present for the other five tasks. The significant correlations between out-of-scanner tests and components 1-3 varied by fMRI task. The first component had the highest correlation with cognitive tests for the Emotional, Gambling, and Relational tasks, while the highest correlations were observed in component 3 for Language and component 2 for Working Memory. We additionally examined correlations with in-scanner performance on the fMRI tasks and the task components (Supplemental Figure 15). There was a general tendency for the strongest correlations between components and performance from the same task, particularly for the Emotional, Language, Relational, and Working Memory tasks. Several correlations were also present across tasks (e.g. Relational component 1 had correlations with Working Memory task performance), which may be explained by correlations across in-scanner task performance or common activity of task positive networks captured by the first component.

**Figure 7:**
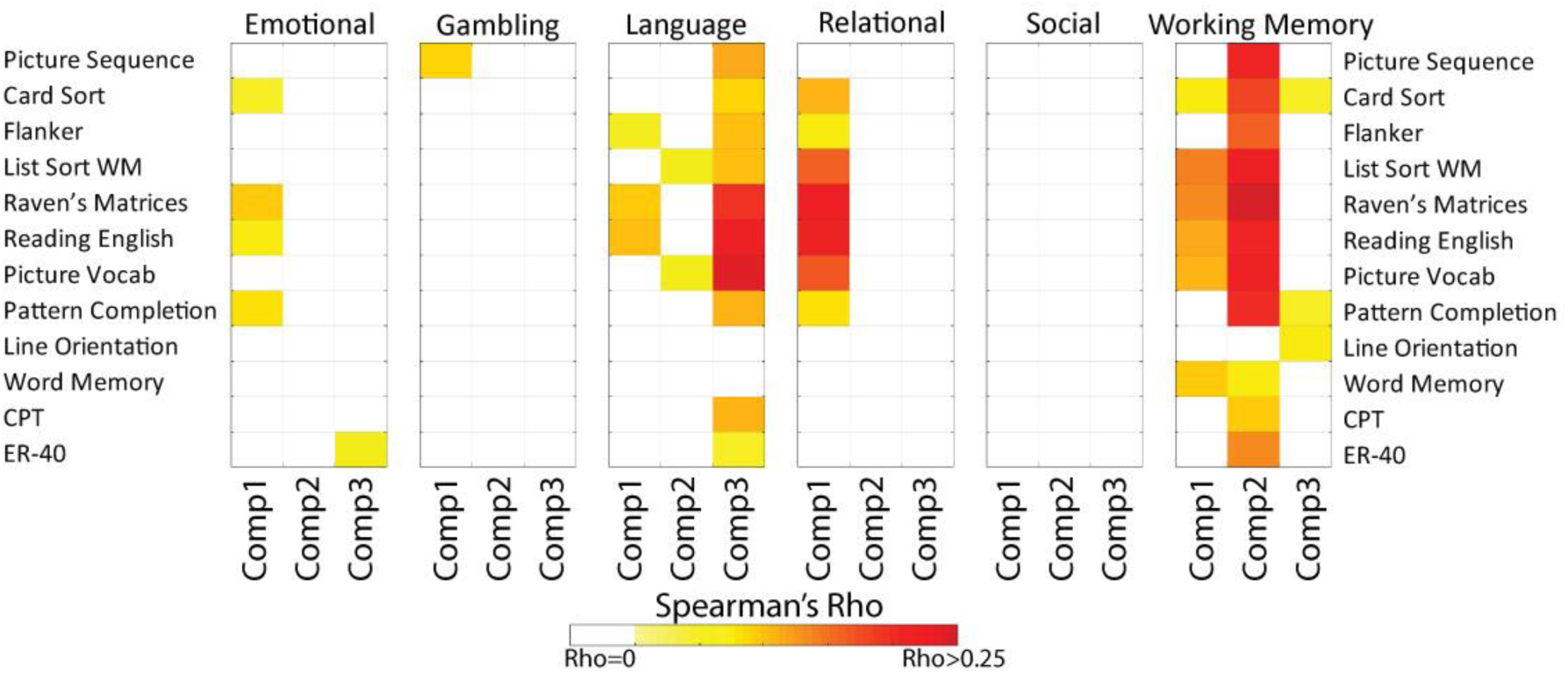
Spearman’s correlations between the first three principal components of the cluster probability matrices and out-of-scanner cognitive scores. For each fMRI task (large boxes) a Spearman’s correlation was calculated for each component (columns labelled as Comp1, Comp2, and Comp3, respectively) extracted from the clustering probability matrix for that task at k=4. Only correlations surviving FDR correction across all presented tests are shown in color (non-significant correlations were set to zero/white). Because the directionality (positive-negative) of the component score is arbitrary, all correlations are presented as positive.

### Pseudo-simulations demonstrate the factors underlying the component score plots

In order to demonstrate that the shape of the component plots were not driven by random noise, a pseudo-simulation approach was used, modifying the input data (t-maps for each participant), and re-performing the bootstrap approach and component plots (Figure 8). First, to consider the effects of global activity on the cluster probability matrices, t-maps for each participant were demeaned, preserving network structure but removing the global component. This resulted in ‘tortilla’ shapes. Second, we randomly shuffled the spatial organization of the t-maps from each participant, removing any cohesive spatial patterns or network structure, while maintaining the mean activity per participant. This transformed ‘tortilla’ shapes into ‘snake’ shapes; these results are consistent with the finding above that the ‘snake’ pattern was associated with differences in mean brain activity. When t-maps were both demeaned and spatially shuffled, the plots collapsed into single points with a small number of random outliers, showing neither a ‘snake’ nor ‘tortilla’ shape emerge from randomized inputs. Finally, the t-maps were randomly shuffled and demeaned, but three simulated ‘networks’ were added, each consisting of approximately 10% of random vertices. Each network was set to a random range of values per participant (i.e. each network could be weakly or strongly ‘activated’ independent of others); one was set to positive, one negative, and the third ranged from negative to positive. Bootstrapping the simulated networks produced ‘tortilla’ shapes, suggesting that the observed ‘tortilla’ patterns arise when multiple brain systems are modulated in a relatively independent way.

**Figure 8:**
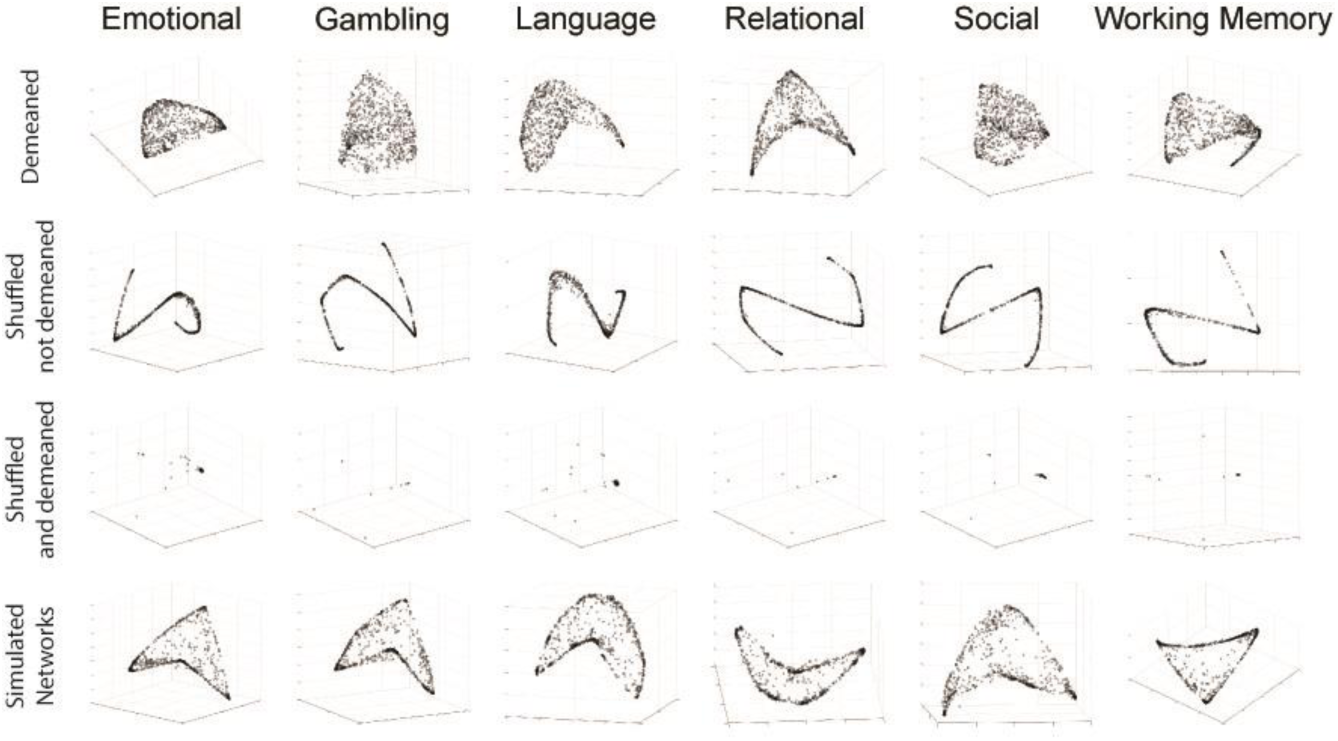
Pseudo-simulations examining the underlying structure of the component space. Cluster bootstrapping was rerun on the originally input t-map matrices after transforms were applied. First, participant t-maps were demeaned prior to clustering, producing ‘tortilla’ shapes (top), then t-maps for each participant was shuffled into a random order but the mean was preserved, creating ‘snakes’ (second row). Data was then both shuffled and demeaned, creating essentially random input, resulting in no shapes (third row). Lastly, data was shuffled and demeaned, and three independent simulated networks’ were introduced; one positive, one negative, and one ranging from positive to negative, recreating the ‘tortilla’ pattern.

### The first component in the Working Memory task does not relate to an abnormal hemodynamic response function (HRF)

Given the global nature of component 1, one possible driving factor is abnormal hemodynamic coupling or a poor fit between the HRF models and individual HRFs. We therefore examined the hemodynamic response function in the Working Memory task. Given the bimodal/anti-normal shape of the distribution, we took the 100 participants with the highest and 100 participants with the lowest component 1 score. The ‘low’ group were those who showed less positive/more negative activity in the 2Back > 0Back contrast. Mean time-series were created for each group (averaging participants and task blocks) to visualize group mean hemodynamic changes during 2Back and 0Back blocks (Figure 9). The morphology of the hemodynamic response was similar between groups. Consistent with the finding of less task-evoked activity in the 2Back > 0Back contrast, the low group showed a reduced response to 2Back and an increased response to 0Back. This suggests that differences in the activity pattern across component 1 represents differences in task-evoked activity, as opposed to abnormal hemodynamic responses.

**Figure 9:**
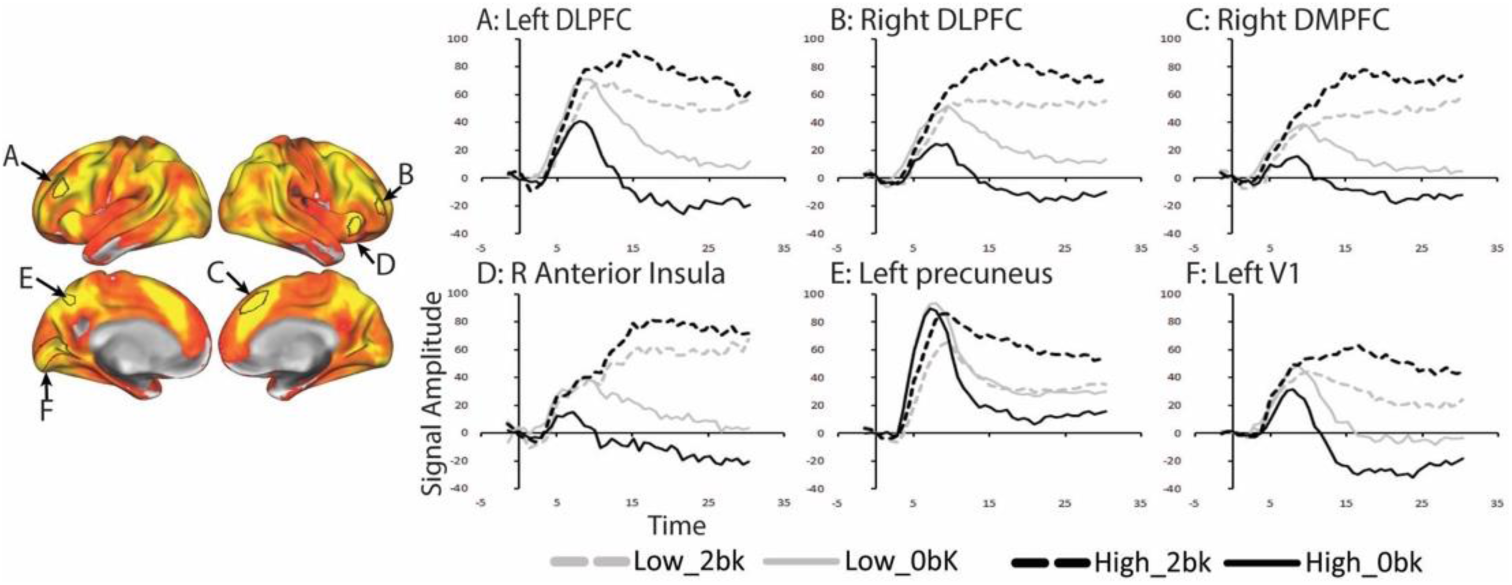
Mean hemodynamic responses to 2Back and 0Back blocks in the Working Memory task for two groups of 100 participants with the highest or lowest scores for the first component of the PCA of the cluster bootstrap matrix for Working Memory (k=4). The ‘low’ group are those with less positive activity in their individual activation maps. ROIs were selected from the Glasser atlas based on the group regression map (left panel).

## Discussion

Standard task fMRI analyses rely on group means to uncover patterns of task-induced neural activity. Our findings show that there is no single ‘group-based’ pattern of activity. The initial clustering solutions found sub-groups of participants along an apparent continuum from ‘deactivators’ to ‘strong/hyper activators’, emphasizing that whole-sample group analyses fail to capture the full range of task-evoked activity^6,29^. Characterizing the range of variability across participants may lead to a more complete understanding of cognitive processing in the brain. The bootstrapping approach allowed a closer examination of the consistency and reliability of the clustering results, while extracting the first three components provided a new approach to quantify the range of variability across individuals. The resultant ‘tortilla’ and ‘snake’ plots represent a lower-dimensional participant similarity space, not only characterizing the range of observed activity but also providing information on the patterns of activity driving this variability. Based on the simulations, the ‘snake’ shape implies a more singular system varying along a continuum, while the ‘tortilla’ shape implies multiple systems which can vary independently. Importantly, both the initial clustering solutions and cluster probability components related to out-of-scanner cognitive performance, demonstrating the behavioral relevance of these components.

Plotting participants in component space quantified the variability in brain activity across fMRI tasks. While some of the patterns of variability were explained by the positive to negative axis (component 1), other important dimensions of variability were also observed, embodied in the ‘tortilla’ and ‘snake’ shapes in the 3D plots of the components; individual variability was represented within a lower multi-dimensional space. This new space might be thought of as a lower-dimensional participant similarity space. While we made use of clustering to define this new lower-dimensional space, the results of this analysis are more in keeping with viewing patterns of activity along continua. The cluster bootstrap approach can be considered an unsupervised feature selection step; across bootstraps participants are clustered according to the most salient features within the data (patterns of brain activity) which define differences and similarities across individuals. It is notable that all tasks fell along one of two spaces: an S-shaped ‘snake’ shape, in which the majority of participants fall along the line with greatest divergence at the curves; and a folded circle ‘tortilla’ shape, in which the majority of participants fell along the circle edge and most/all participants were on the ‘surface’. Simulations demonstrated that the ‘snake’ pattern was related to some variance of mean activity (though this may be driven by greater activity within specific task relevant systems), while the ‘tortilla’ shape appears to be a result of multiple systems, which can potentially independently vary in amplitude. This new participant similarity space represents a manifold demonstrating the range of patterns of activity during a cognitive task, which may have important behavioral or clinical implications for mapping brain-behavior relationships.

Consistent with the initial cluster solution, we found relatively global patterns of activity related to component 1, suggesting a positive to negative ‘axis’ of activity across participants. Prior work focused on task connectivity has suggested an increase in global integration during task performance^30,31^ which may in part explain why such a wide range of brain regions were related to component 1. While variability across fMRI task activity patterns have been highlighted by previous studies^4,6^, to our knowledge this pattern of activity ranging from positive to negative across participants has only been suggested in our previous single-task study^3^. At a network level, the first component consistently had high loading on the dorsal attention network, which is related to task-oriented top down attentional processing^32,33^, and serves a more domain-general rather than task-specific role in cognitive processing^28,34-36^. Although this could indicate that clustering is driven by attention, reductions or variance in attention would be expected to have behavioral consequences in task performance, and the first component was not strongly related to in-scanner task performance in most cases (Supplemental Figure 16). This argues against attention as the primary factor underlying the first component, although attention likely represents some of the observed variance.

We favor a behavioral explanation as the primary driver of cross-task variability: the pattern observed in a given participant is strongly influenced by the specific cognitive process or strategy they are using during the task. Previous work has shown how different cognitive strategies can modulate brain activity^6-8,37,38^ and that training participants to make use of alternate strategies can change observed patterns of activity^9,39^. This was supported by the lack of overlap we found between clustering solutions across tasks. Cognitive tasks can be performed via different operational cognitive procedures (cognitive strategies), which can modify brain activity^7,37,40^. For example, one strategy during the N-Back might be first updating the memory set to include the current and past two items and then matching the first and last items in the set, while an alternative strategy might recall the previous two items as part of an encoded set, then matching the current item and updating the memory set. Participants with fewer cognitive resources might instead utilize a less cognitively intensive approach to matching (if an item feels familiar/recent mark it as a match)^41^, which may not require specifically maintaining and updating the memory set. Presumably this latter strategy would be less efficient in terms of accuracy, but is a viable approach that would make use of alternate brain systems. We hypothesize that this strategy would produce less activity regions associated with working memory (e.g. the fronto-parietal network) and poor task performance, such as was seen in the non-activating Working Memory subgroup. However, caution is required when making inferences on the underlying cognitive strategy based on group maps^29^. Varying cognitive strategies could also explain why component scores were not correlated across tasks: specific cognitive approaches may not transfer across tasks^41^. Effort may also play a role in behavioral variability, related to the importance of the dorsal attention network in the first component.

An alternative explanation for the observed clustering is grounded in the underlying biology of participants; the pattern of activity across tasks is mainly related to the functional strength of those brain systems within those participants. Information flow across regions during task performance has been shown to follow underlying resting state network architecture^24^ which is stable across cognitive tasks^42^ and resting state networks have been used to predict patterns of task induced activity^22,43^. This would suggest that the underlying network function plays a role in task activity, which in turn might indicate that participants with similar network structure might show similar activity patterns across tasks. While we did not find common clustering across tasks, network function likely plays a role in task activation patterns and could explain some of the individual variability seen in patterns of task activity. For example, Gratton and colleagues^42^ examined repeated tasks within a small highly sampled set of individuals and found that group network architecture and inter-subject variability explained important variance in network connectivity. A task by individual interaction explained some variance in connectivity profiles, while task alone explained little variance^42^. This suggests that the manner in which participants individually modulated their functional networks during tasks was more relevant than the task itself. The individual variability is more likely to represent a trait effect, while the individual-by-task variability is more likely to reflect individualized state changes related to task demands. This highlights the importance of both the underlying network structure (a biological factor) and individually specific patterns (potentially related to different cognitive task approaches) in task connectivity.

Our current approach, and task analyses in general, do not consider changing neural dynamics during task performance^30,31,44^. Our analysis reveals variation in brain activity *among* individuals, but does not account for variability across trials *within* an individual. Thus, variation in brain activity may reflect the portion of time occupying different and fluctuating brain states during the course of a task^30^. Future work may incorporate more dynamic information to better understand how fluctuating neural states may relate to individual patterns of activity across the whole task.

These findings highlight the importance of individual variability when considering patterns of task-related brain activity related to cognitive processes. While group maps may be informative, they fail to capture the range of variability present across the population. Individuals along the more extreme ends of the spectrum in terms of brain activity are still performing the cognitive functions being measured in a given fMRI task, though their individual brain pattern may diverge very substantially from the average map. Understanding the range of this variability can help us better understand the complex neurobiology underlying cognitive processing, with implications for both understanding cognitive systems as well as for clinical research comparing activity across groups.

## Methods

### Participants

Data were obtained from the HCP S900 release^45,46,20^, ranging in age from 22 to 35. Of 899 participants with fMRI data available, only those who completed all six fMRI cognitive tasks (Emotional, Gambling, Language, Relational, Social, and Working Memory) were included, leaving a sample of 822.

### Subject-level activation maps

Task activation maps from specific contrasts across six cognitive paradigms in HCP were used, including (selected contrast in parentheses): Emotional (Faces-Shapes), Gambling (Reward-Punish), Language (Story-Math), Relational (Related-Match), Social (TOM-Random), and N-Back Working Memory (2Back-0Back). Individual subject statistical analyses provided as part of the HCP S900 release^20,47^ were used for all further analysis. To facilitate group-wise comparisons, data were selected from the surface smoothed (8mm FWHM) using the HCP minimal preprocessing pipeline^47^. This pipeline includes motion correction, distortion correction, registration to standard space, and generation of a grayordinate (cortical surface) time-series for each task run. For each task, the HCP collected two runs which differed with respect to the MRI phase encoding direction (left to right, or right to left). Statistical analyses were performed in FSL. Fixed-effects (“first-level”) analysis was performed on each run separately, including the smoothing stage (performed on the cortical surface), and then the two runs for each task were combined via a second fixed-effects analysis. For each participant, the HCP provided t-maps for each task were used. We used t-maps as opposed to Beta estimates for cluster analysis as the t-values are deweighted in noisy voxels due to the higher variance.

### Initial cluster analysis

All clustering analyses were completed in MATLAB (R2017b). Hierarchical clustering with Euclidean distance and Ward’s linkage was used ^3,21^. In order to perform the clustering, a ‘spatial’ matrix was created by extracting the t-statistics across the cortex for each participant as a vector and stacking participants (producing a matrix of spatial activation across subjects). Clustering was then run on each task separately, producing solutions from k=2 to k=10 clusters. An additional analysis was run using K-means, with k=4, for visual comparisons with the results from hierarchical clustering.

### Group analysis by cluster

In order to visualize patterns of activity within each cluster, group-level maps were created for each separate cluster. Second-level (group) one-sample t-tests were performed using SPM12, using the contrast maps provided by HCP for the contrast of interest. The statistical threshold for the group/cluster map is dependent upon sample size. That is, a smaller sample requires a larger t-value to establish significance. In order to allow for a clearer visual comparison between groups, a minimum t-value was set of t=5, which approaches or exceeds p<0.05 FWE at the single vertex level for most clusters (if n ∼>100). By comparison, p<0.001 uncorrected t-values ranges from about 2.8 to 3.2 for most cluster sizes. Thus, this t-value avoids individual thresholding for each cluster solution (which may make direct visual comparisons across maps challenging) but is a stringent correction for multiple comparisons^48^.

### Cognitive data

Cognitive data consisted of 12 measures available battery within the HCP protocol. From the NIH toolbox, age-adjusted scores were taken from the Picture Sequence test (memory), List Sort test (working memory), Dimensional Card Sort test (executive function, cognitive flexibility), Flanker task (inhibition), the Pattern Completion test (processing speed), Reading in English test, and the Picture Vocabulary test. Additional cognitive measures included were the Short Penn Continuous Performance task true positives (sustained attention), Penn Word Memory Test (verbal episodic memory), the Variable Short Penn Line Orientation test median correct reaction time (spatial orientation), Raven’s Progressive Matrices correct responses (fluid intelligence), and the Penn State Emotional Recognition test reaction time (emotional processing). All cognitive scores were Z-transformed (within the sample) to a mean of zero and standard deviation of one to allow direct comparison across tests. Initially, a PCA was run on all 12 test scores for data reduction purposes; however, the first three components only accounted for 25%, 15%, and 9% of the variance, respectively. Therefore, all 12 scores were used in subsequent analyses. Selected relevant within-scanner fMRI task performance variables were also extracted for each of the six fMRI tasks (24 scores total), and Z-transformed to a mean of 0 and standard deviation of 1.

Differences in cognitive scores between clusters were performed across all cluster solutions (k=2 to k=10) for all six tasks for all 12 scores (i.e. a total of 648 one-way ANOVAs were performed), to examine differences in cognitive scores between clusters. Significance was defined as p < 0.05, FDR corrected across all 648 tests performed.

### Cluster bootstrap analysis

The cluster bootstrapping analysis was performed via a bootstrapping without replacement approach. Bootstrapping was run for each of the six tasks and each cluster size from k=2 to k=10. For each task at each value of k, 1000 permutations were performed, building a cluster solution using a random 75% of the sample. For each combination of task and k value (e.g. Working Memory task, k=4), a ‘clustering probability’ matrix was calculated to capture the probability of each pair of participants clustering together. This was quantified as the probability (from 0 to 1) of a pair of participants being in the same cluster when they were both in a bootstrap. This resulted in an NxN matrix of clustering probabilities. For visualization purposes, an optimal ordering function (MATLAB’s optimalleaforder.m) was run using hierarchical clustering with Euclidean distance and average linkage to sort the matrix in such a way as to maximize the sum of the similarity between adjacent participants. This was used for sorting/ordering purposes only and resulted in a more visually informative way by placing similar participants together.

### Bootstrap components analysis

Examination of the agreement matrices indicated that the data did not necessarily fall into distinct clusters, but a large proportion of variance was represented by the position on the diagonal of the matrix. However, the matrices also suggested there may be more than a single dimension to the data (i.e. off-diagonal ‘bulges’). In order to better understand the factors driving clustering results, a Principal Component Analysis (PCA) was run in MATLAB on the agreement matrices to extract the top three components. Note that these components were not taken from the t-maps, but from the cluster bootstrap matrices. The components are therefore representations of the lower-dimensional factors which cause participants to cluster together or not. Data were visualized by plotting participants into 3D space using the component scores for each participant.

To assess behavioral relevance, a Spearman’s correlation was conducted between the component scores and 12 cognitive scores (the components did not follow a normal distribution, see Supplemental Figure 15). The top three component scores for k=4 for each task were compared to each of the 12 cognitive scores (3 scores × 12 tests × 6 tasks = 216 total correlations); significance was determined by FDR correction applied to the full set of 216 correlations.

### Regressing PCA scores to recover associated patterns of brain activity

In order to map the patterns of brain activity underlying the PCA scores, a second level analysis of brain activity was run in SPM12, using the top three component scores as regressors of interest. This group-level analysis identified specific brain regions in which participants with higher component scores showed greater or lower levels of task evoked activity, and thus the regions which were related to the component scores.

### Pseudo-simulations reproducing the component score plots

In order to identify the characteristics of variations in functional activity driving the ‘tortilla’ and ‘snake’ shapes in the cluster probability component plots, a pseudo-simulation approach was undertaken. The fMRI t-maps which were used in the bootstrap analysis were used, but modified to explore different hypotheses; 1) cluster probability may be related to mean brain activity across the cortex; 2) cluster probability may be related to activity in independent systems; or 3) cluster probability may be an element of random similarities between scans. Pseudo-simulations were conducted by performing specific operations on the input data matrix (participant’s t-statistics cortical vertices; t-maps), rerunning the cluster bootstrap across 1000 iterations at k=4 for each task, calculating the first three principal components of the pseudo-simulated cluster probability matrix, and plotting each participant in 3D component space. Modifying the actual t-maps allowed us to consider the effects of different aspects of the data which may underlie the observed cluster shapes on simulated data with similar characteristics/distribution to the original data. In the first simulation, t-maps from each participant were demeaned; this leaves relative differences across brain networks while removing any potential effects of overall activity across the brain. In the second simulation, the spatial order of vertices for each participant were randomized, removing any coherent spatial organization of activity between participants, but the mean was maintained. This examined the effects of mean differences in brain activity between participants when no network structure was commonly present between participants. In the third simulation, t-maps were spatially shuffled and demeaned; this examined the effects of removing spatial relationships across individuals, while maintaining a realistic distribution of t-statistics within individuals. Finally, for the forth simulation t-maps were demeaned and shuffled, but three simulated networks were added to the data, each independently modulated by creating network weights; ranging from −1 to 1 (for a network ranging from negative to positive), from 0.0012 (1/822; multiplying network by zero would produce unrealistic results) to 1 (for a positive weighted network) and − 0.0012 to −1 (for a negative weighted network). Each network had 822 unique weights (1 per individual), which were randomly assigned to participants, randomly across networks. As such, each network was independently modulated across participants (e.g. it was possible to have high positive and low negative network modulation, or low both, high both, or one intermediate and the other high or low, etc.). This pseudo-simulation tested the effects of independent networks which could be more or less engaged across participants, causing both localized differences as well as a distribution of mean activity.

### Hemodynamic response related to the first component in Working Memory

Given the bimodal/anti-normal shape of the distribution for component 1 of the clustering probability matrix in working memory, we took 100 participants from each end of the distribution (i.e. the 100 participants with the highest and 100 participants with the lowest component score). The ‘Low’ group were those who showed less positive, more negative activity in the 2Back > 0Back contrast. Time-series regions of interest (ROIs) were extracted from ROIs from the Glasser HCP parcellation^49^ from the preprocessed but unsmoothed time-series (dtseries) fMRI files. For a selected group of ROIs, we extracted the time-series and created an average of 0Back and 2Back blocks (e.g. the volumes corresponding to the task blocks), including two volumes before task onset and lasting 30 seconds (the task block duration). Mean time-series were created for each group (averaging participants and task blocks) to visualize group mean hemodynamic changes during 2Back and 0Back N-Back performance. Mean time-series were baseline corrected to a baseline mean of zero using the first 5 volumes (2 before and 3 after task onset).

## Code Accessibility

Code and processed data is made available via github (https://github.com/colinhawco/HCP_cluster_analysis).

## Acknowledgements

CH was funded through the Brain and Behavior Research Foundation. The authors would like to thank Dr. Randy McIntosh for helpful comments on an earlier version of this manuscript, and Joseph Viviano for technical help and comments.

## Author Contributions

CH led the design, analysis, impetration, and writing of the manuscript. EWD provided scientific ideas, interpretation, and analytical support. GJ contributed to drafting and revising the manuscript and figures, and interpretation of results. ZJD provided scientific contributions and overall supervision. AV contributed to methods and interoperation and well as providing overall supervision and access to computational resources. All authors contributed to the final manuscript.

