## Supplementary_Figures for "Moving Beyond the Mean: Subgroups and dimensions of brain activity and cognitive performance across domains"

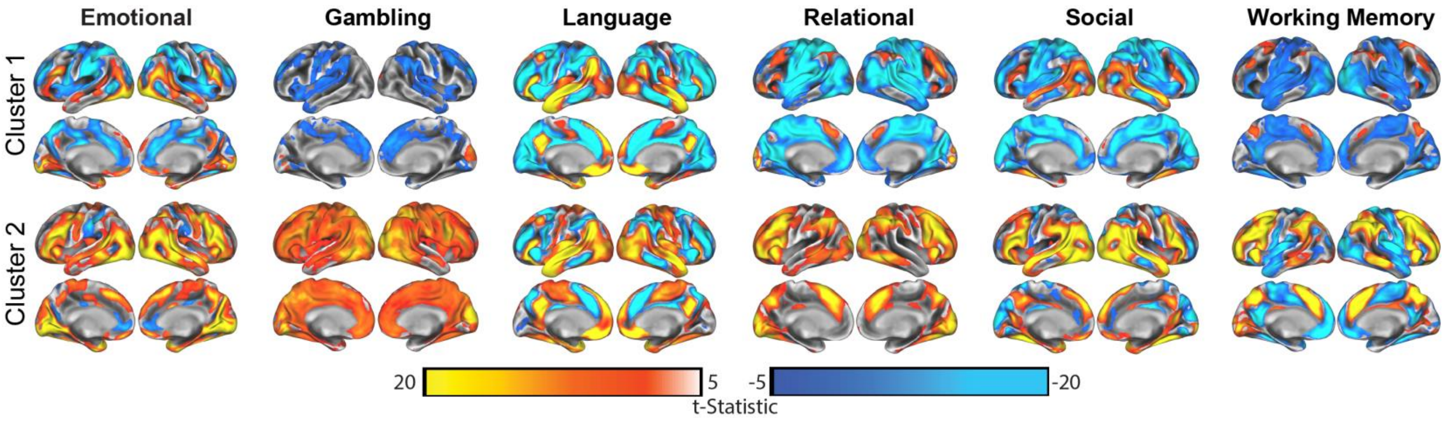


**Supplemental Figure 1**: Group analyses for each task (columns) by cluster (rows) for k=2. All maps are thresholded at t>=5 for visual comparisons. Group analyses (one-sample t-tests) were completed in SPM12. The left hemisphere is shown on the left for each image.

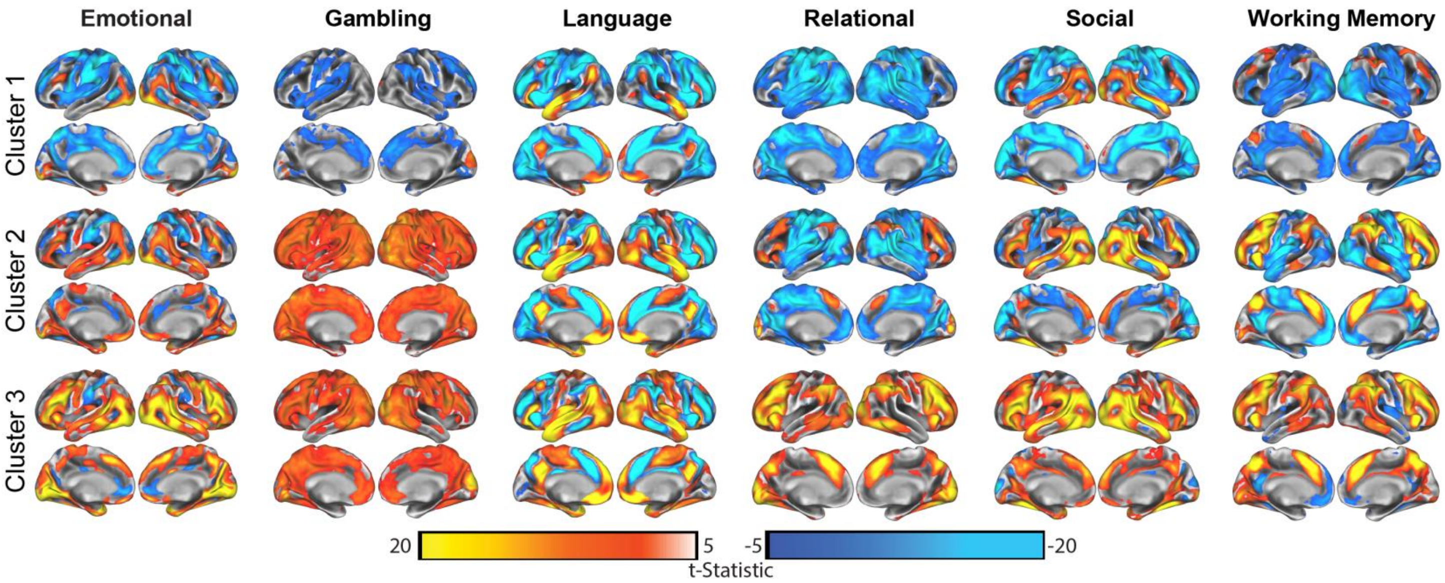


**Supplemental Figure 2**: Group analyses for each task (columns) by cluster (rows) for k=3. All maps are thresholded at t>=5 for visual comparisons. Group analyses (one-sample t-tests) were completed in SPM12. The left hemisphere is shown on the left for each image.

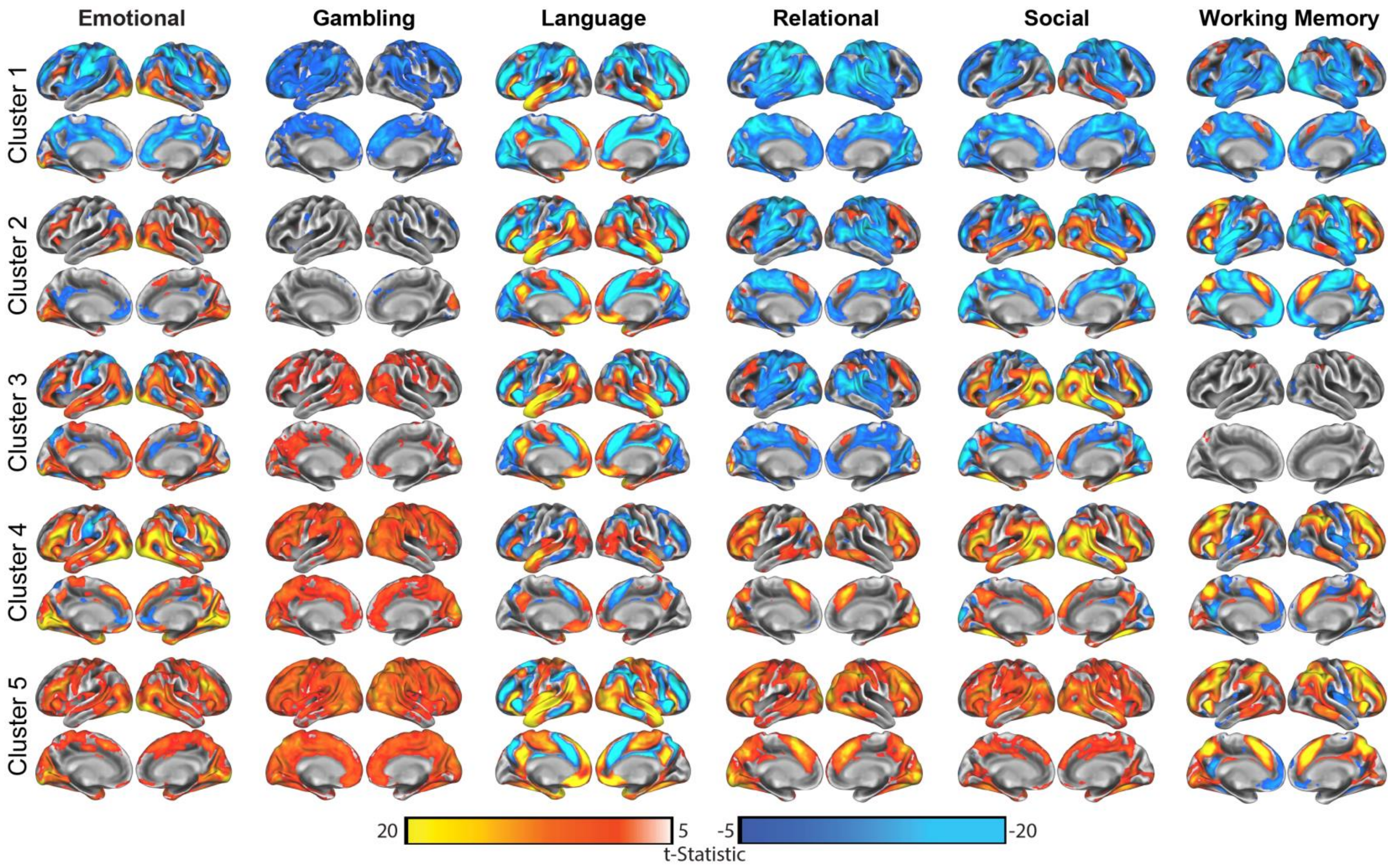


**Supplemental Figure 3**: Group analyses for each task (columns) by cluster (rows) for k=5. All maps are thresholded at t>=5 for visual comparisons. Group analyses (one-sample t-tests) were completed in SPM12. The left hemisphere is shown on the left for each image.

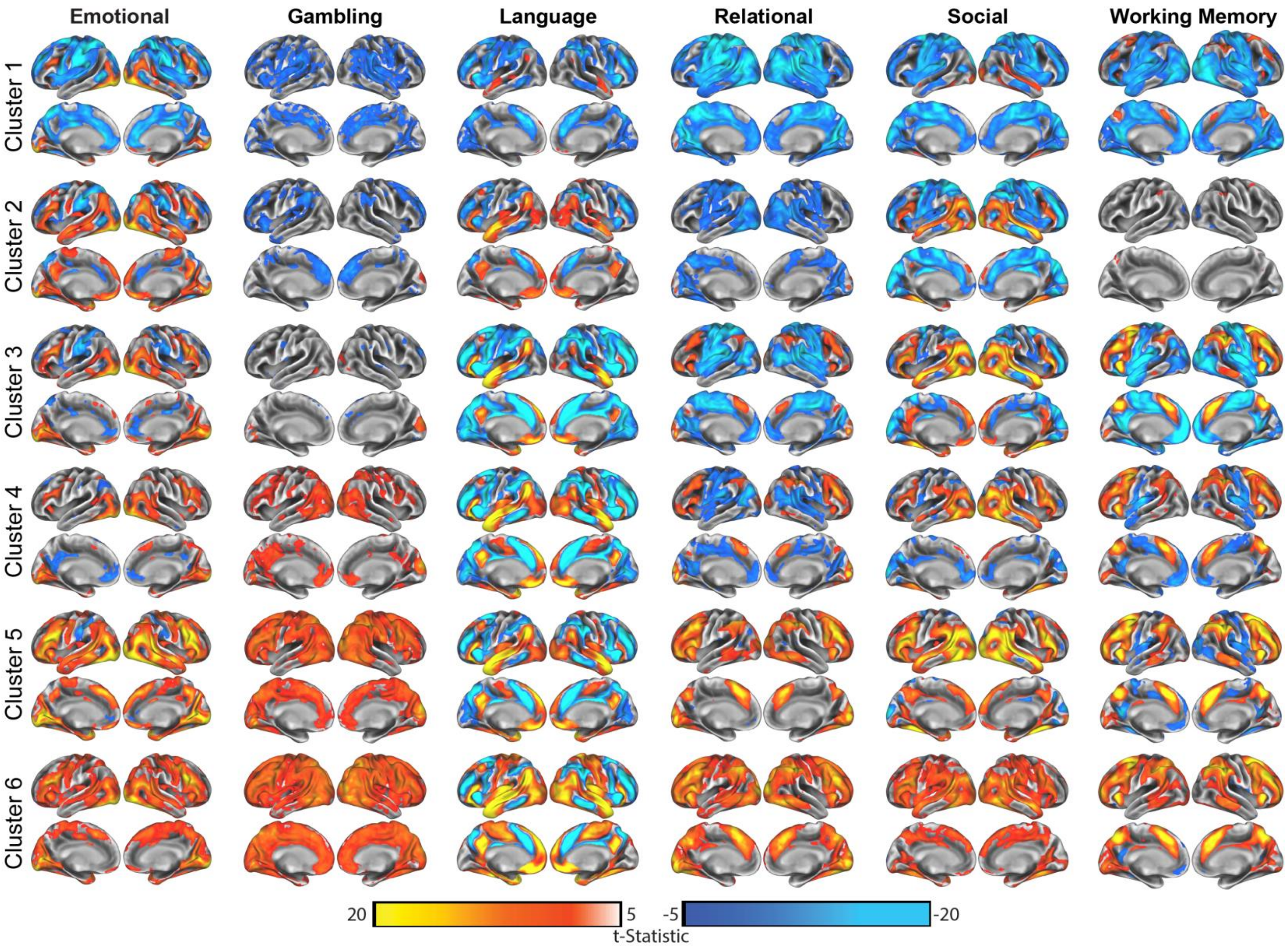


**Supplemental Figure 4**: Group analyses for each task (columns) by cluster (rows) for k=6. All maps are thresholded at t>=5 for visual comparisons. Group analyses (one-sample t-tests) were completed in SPM12. The left hemisphere is shown on the left for each image.

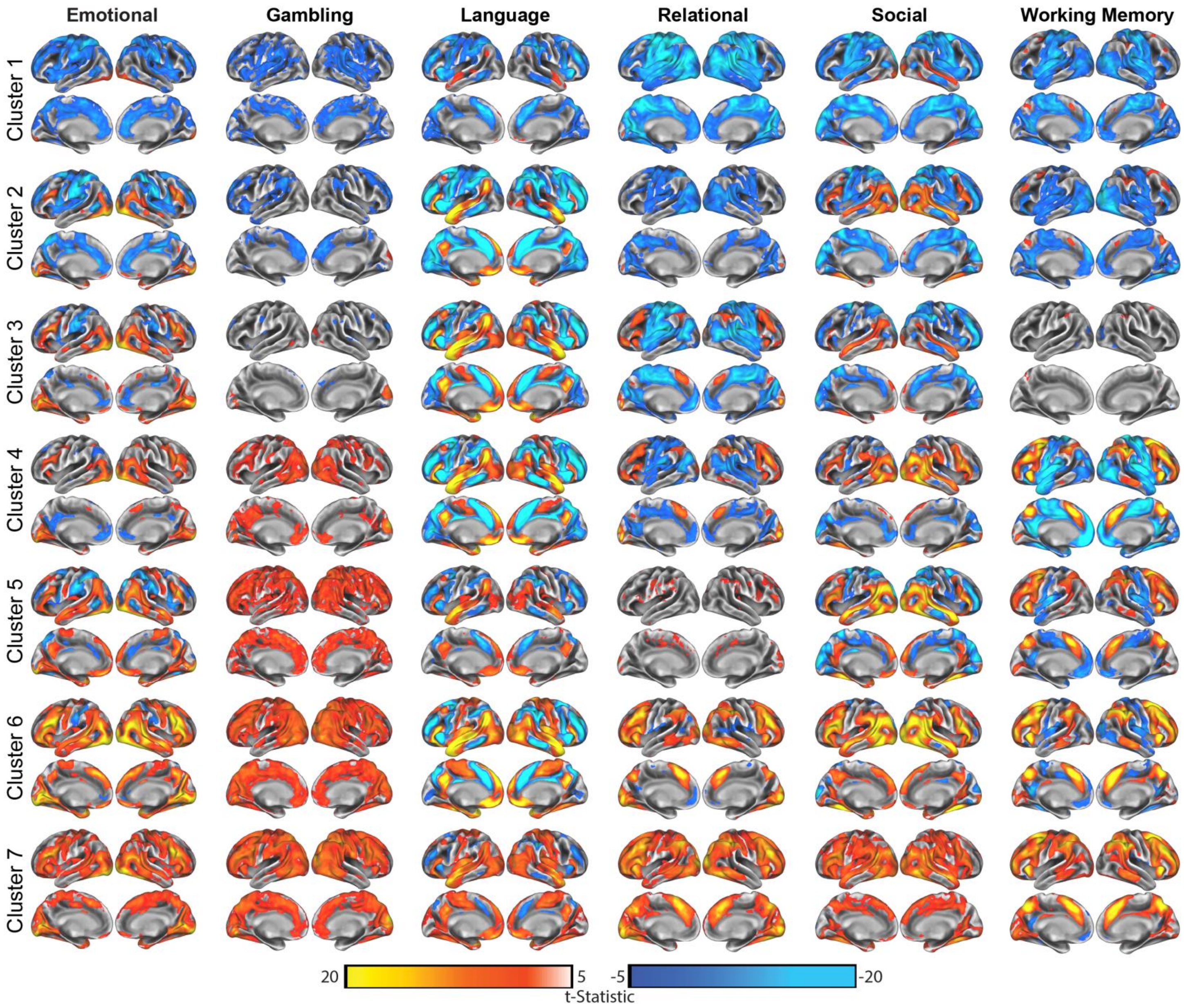


**Supplemental Figure 5**: Group analyses for each task (columns) by cluster (rows) for k=7. All maps are thresholded at t>=5 for visual comparisons. Group analyses (one-sample t-tests) were completed in SPM12. The left hemisphere is shown on the left for each image.

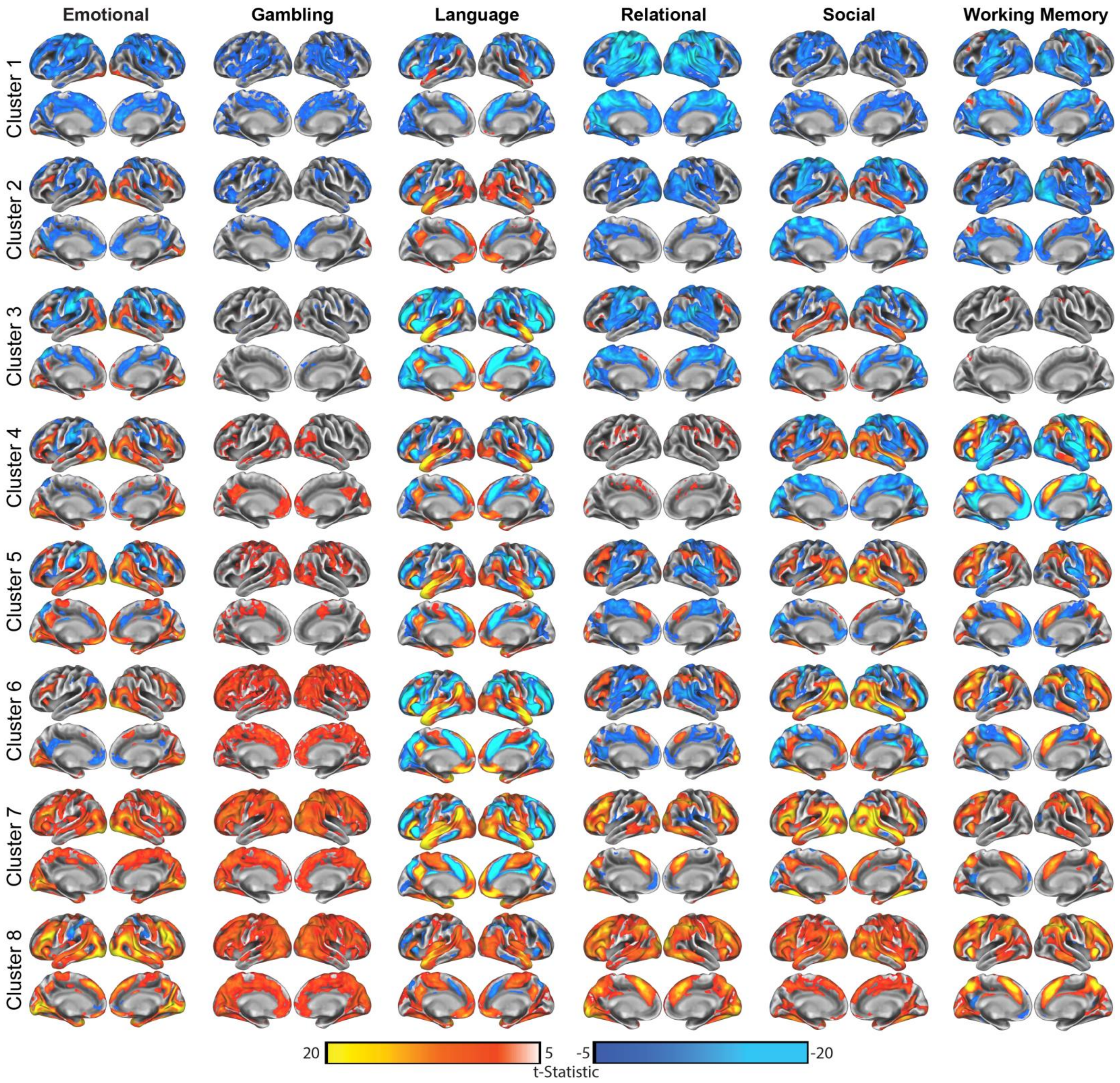


**Supplemental Figure 6**: Group analyses for each task (columns) by cluster (rows) for k=8. All maps are thresholded at t>=5 for visual comparisons. Group analyses (one-sample t-tests) were completed in SPM12. The left hemisphere is shown on the left for each image.

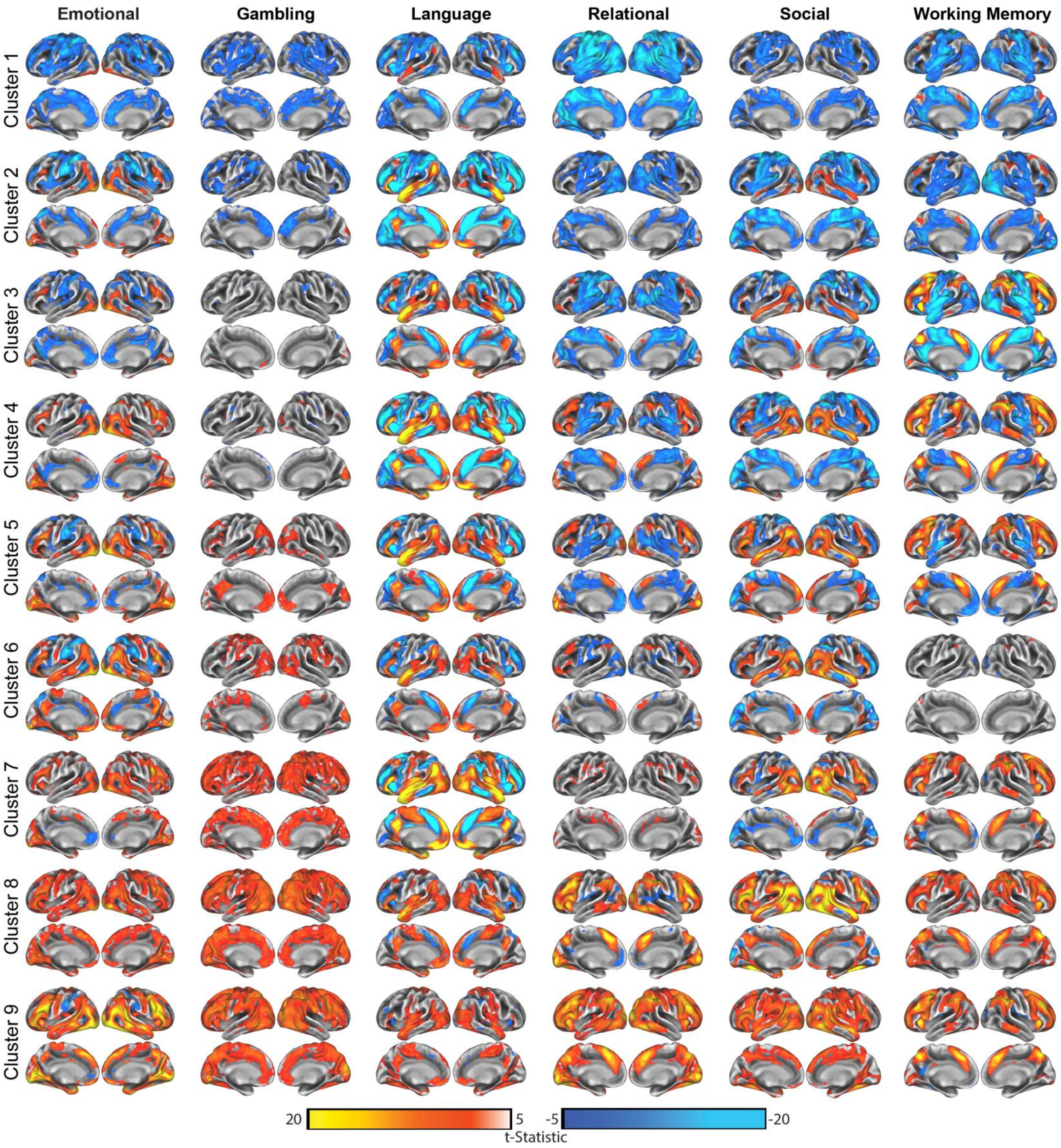


**Supplemental Figure 7**: Group analyses for each task (columns) by cluster (rows) for k=9. All maps are thresholded at t>=5 for visual comparisons. Group analyses (one-sample t-tests) were completed in SPM12. The left hemisphere is shown on the left for each image.

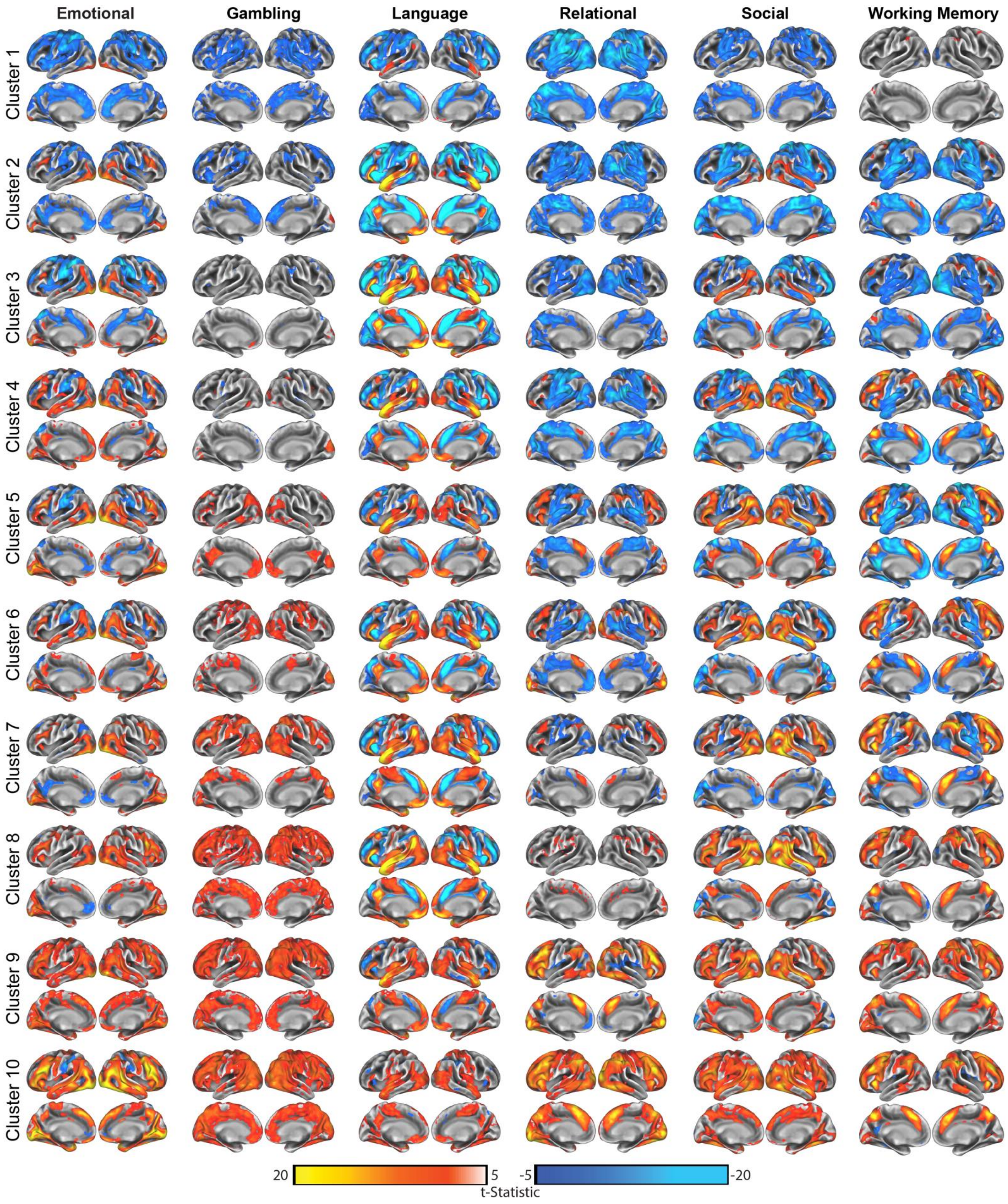


**Supplemental Figure 8**: Group analyses for each task (columns) by cluster (rows) for k=10. All maps are thresholded at t>=5 for visual comparisons. Group analyses (one-sample t-tests) were completed in SPM12. The left hemisphere is shown on the left for each image.

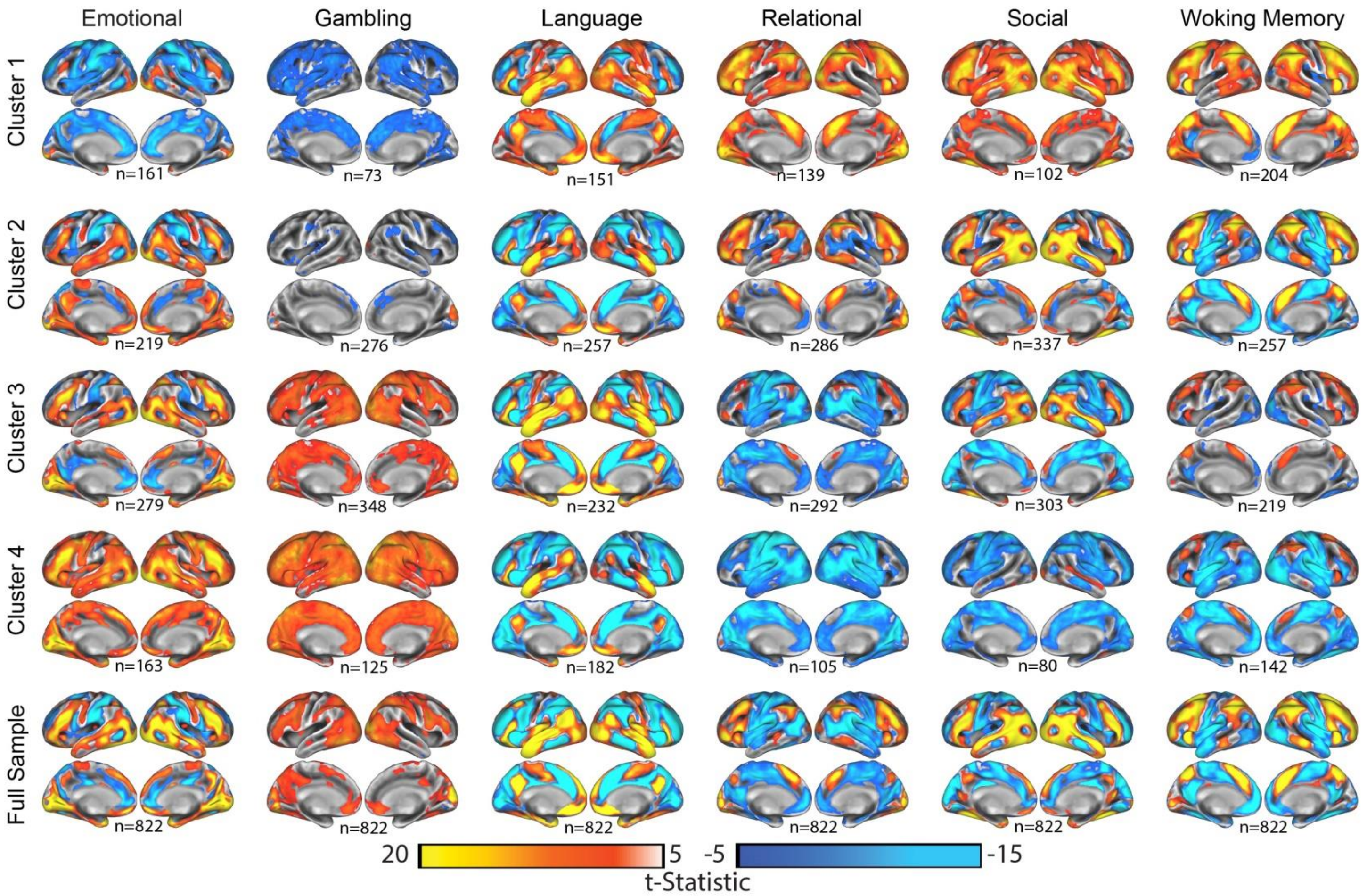


**Supplemental Figure 9**: Group statistical maps across clusters when K-means clustering was used (k=4). Similar to the results observed with hierarchical clustering, the dominant pattern is a range from predominantly negative activity to predominantly positive activity.

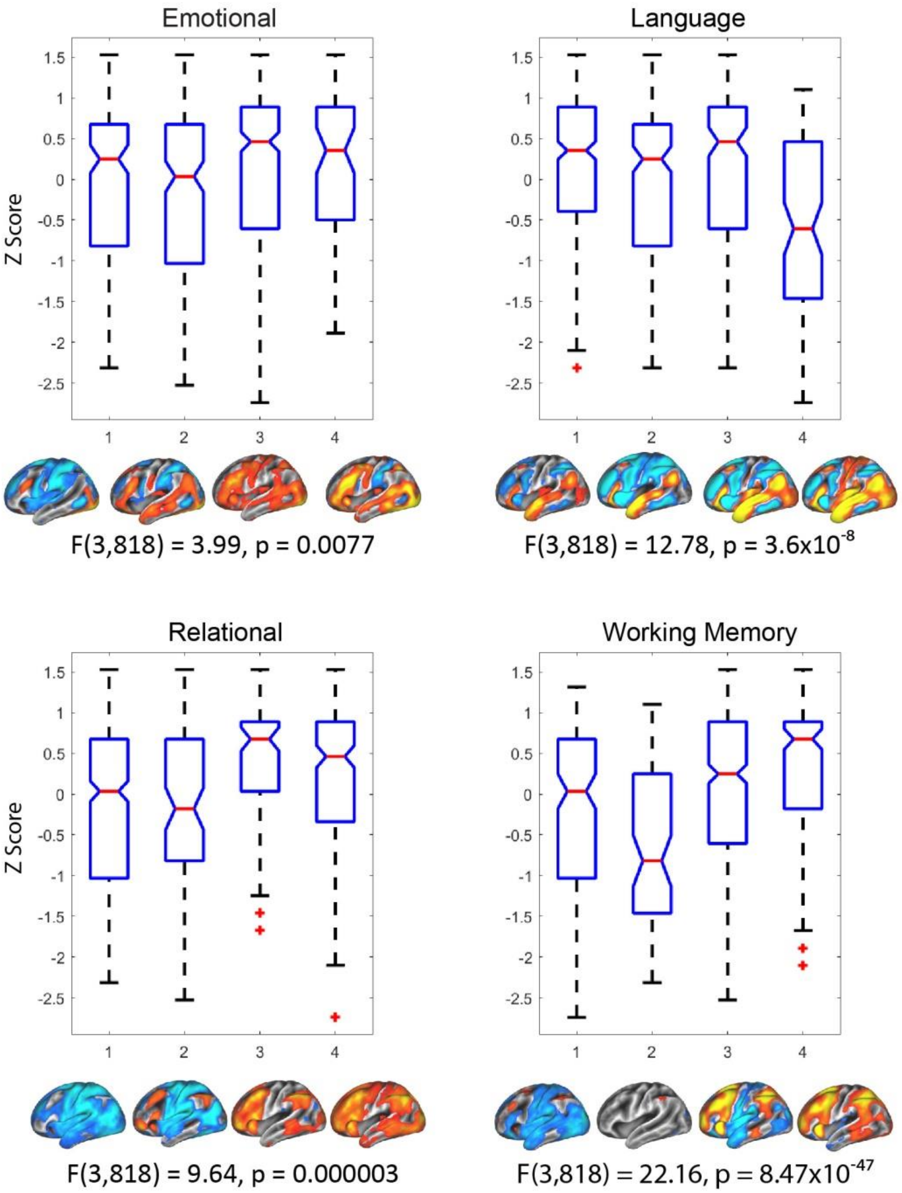


**Supplemental Figure 10**: Z-transformed scores on Raven's Progressive Matrixes (PMAT) for the four tasks showing PMAT differences between clusters. Clusters are arranged as in Figure 1 (deactivating cluster on the left, strongest activating cluster on the right), with left lateral hemisphere activity shown for reference. Boxplots show 25th to 75th percentile of scores, whiskers include non-outlier points, while outliers are marked via a cross. F statistics for the one-way ANOVA for cluster differences are shown below each boxplot.

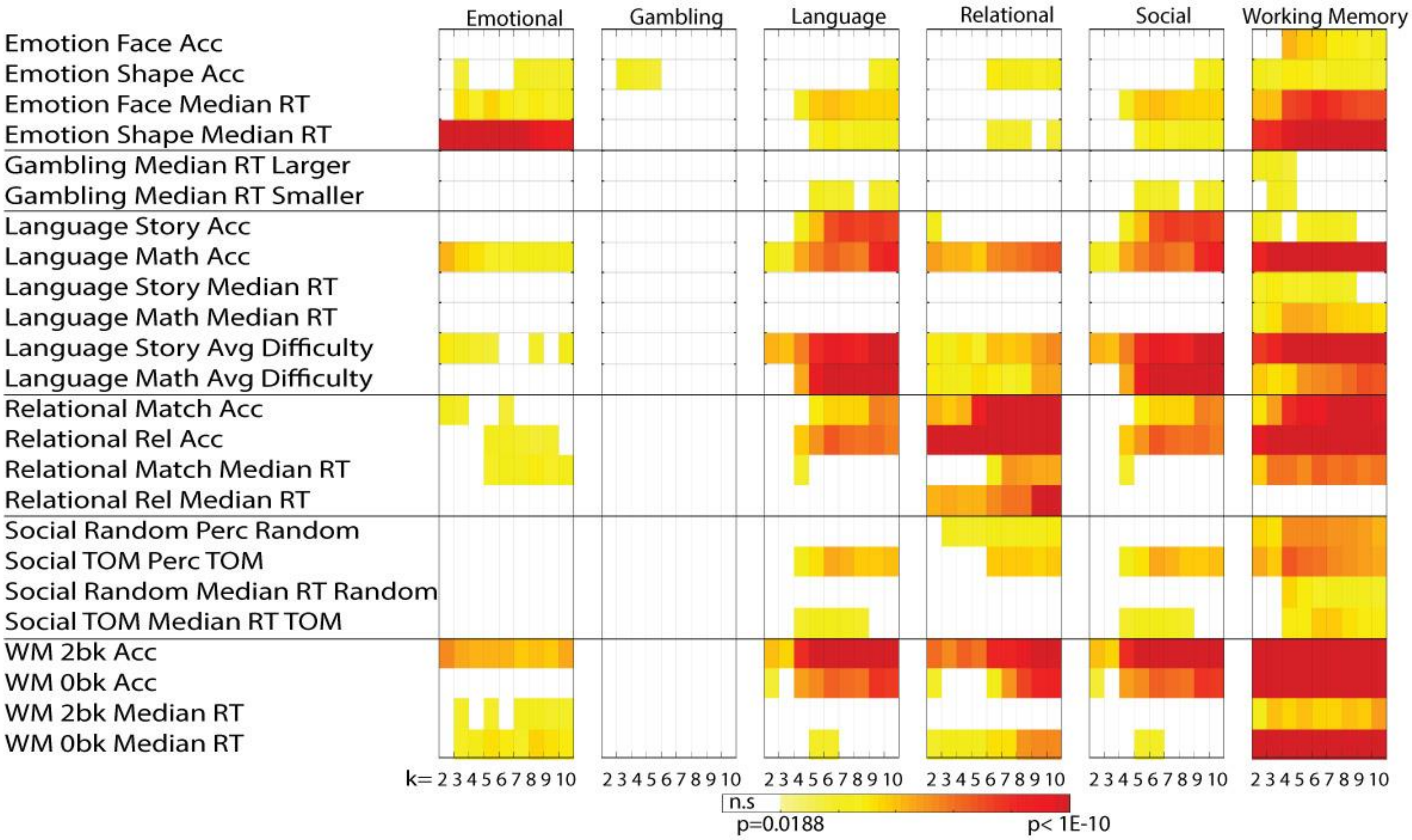


**Supplemental Figure 11**: Results of one-way ANOVAs comparing differences between clusters on performance during the fMRI tasks. For each fMRI task (large boxes) a one-way ANOVA was run for each cluster solution (k=2 to k=10; columns inside boxes) for each cognitive test (rows within boxes). P-values from each ANOVA are presented as a colored box. ANOVAS were run for the initial clustering solution using hierarchical clustering. Any p-values which were not significant (FDR corrected across all tests, p<0.05) are shown as white.

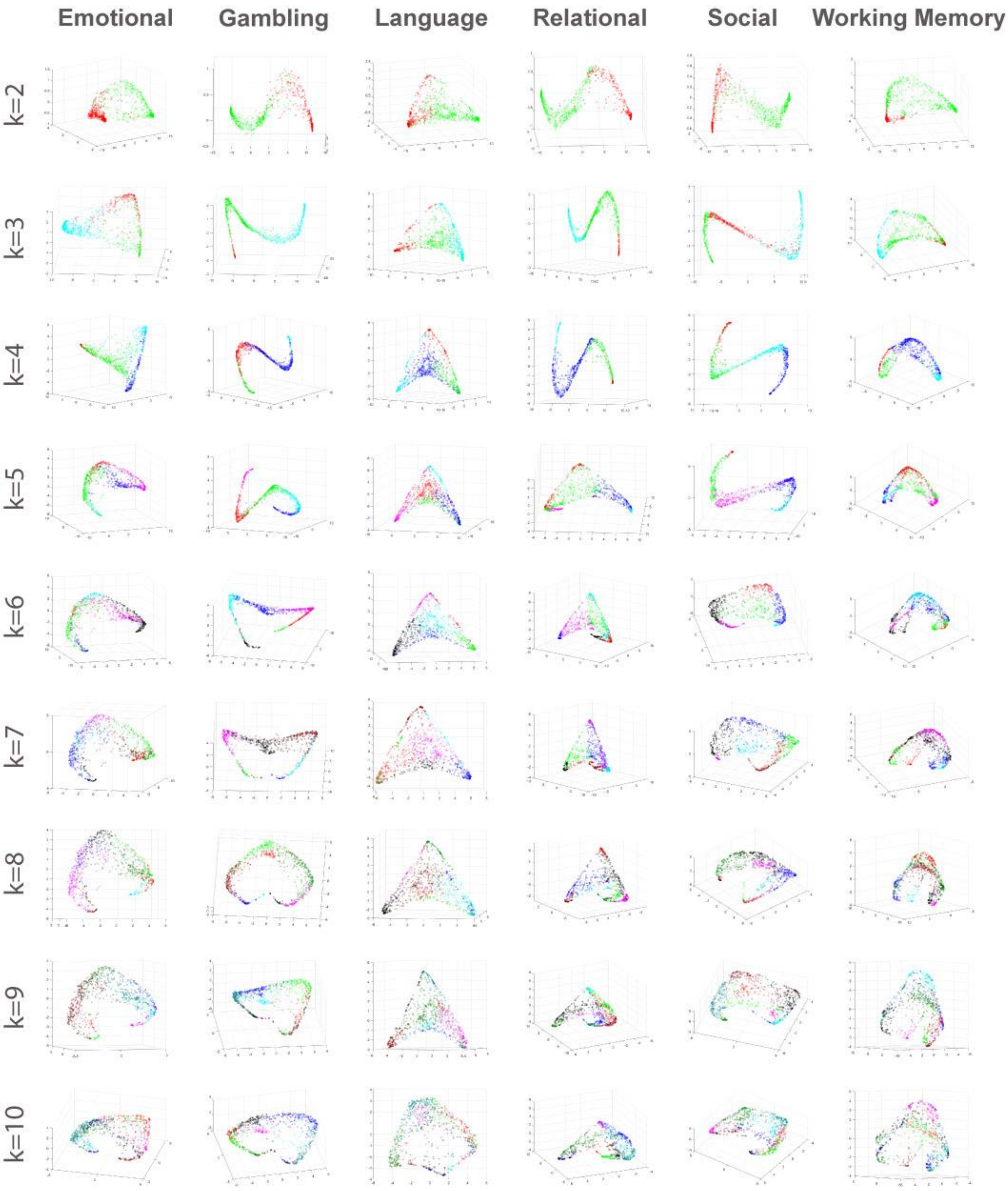


**Supplemental Figure 12**: Participants mapped into 3D space based on their scores in the first three components of the cluster probability matrix, for all values of k in all tasks. Colors represent cluster membership in the whole sample clustering solution (as in Figure 1 and Supplemental Figures 1-8). While Relational and Social show 'snake' shapes at k=3 and k=4, they merge towards a 'tortilla' shape at higher values of k. Additionally, at higher values of k, the manifold becomes less cohesive.

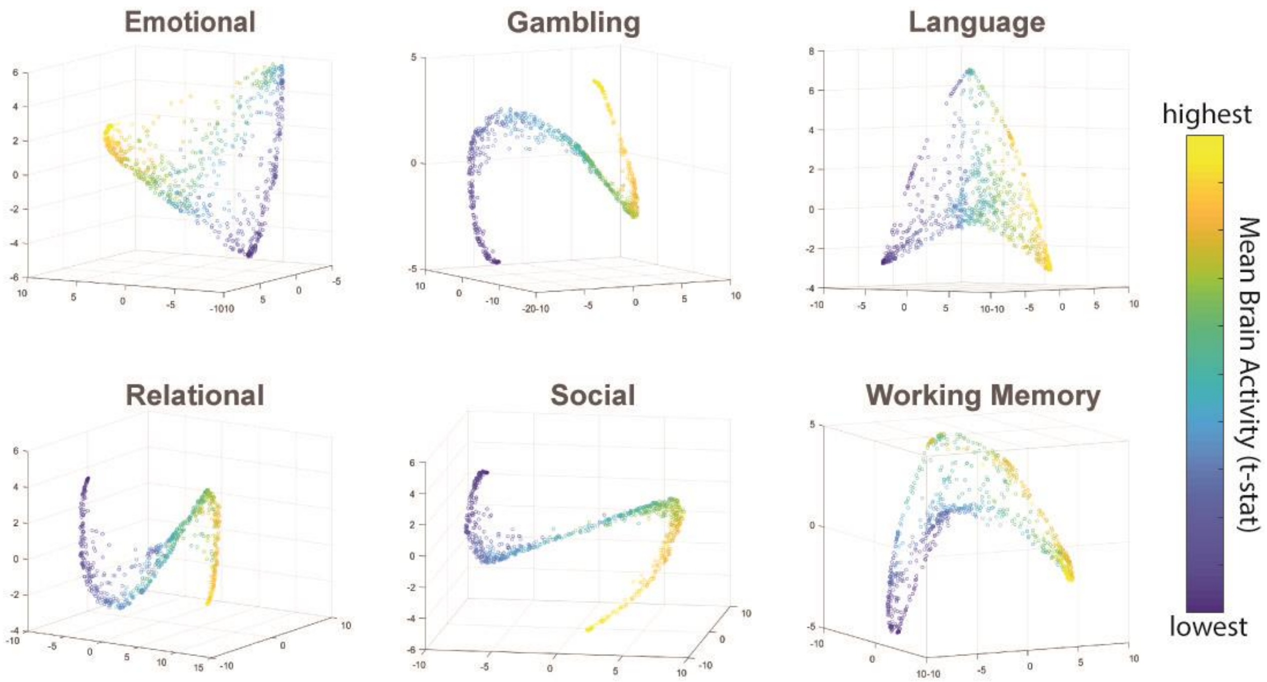


**Supplemental Figure 13**: Participants mapped into 3D space based on their scores in the first three components of the cluster probability matrix (k=4), colored along the Parula spectrum (color bar) by mean brain activity. Mean activity was defined as the mean of all t-stats within the cortex for each participant for the contrast map used in the clustering analysis.

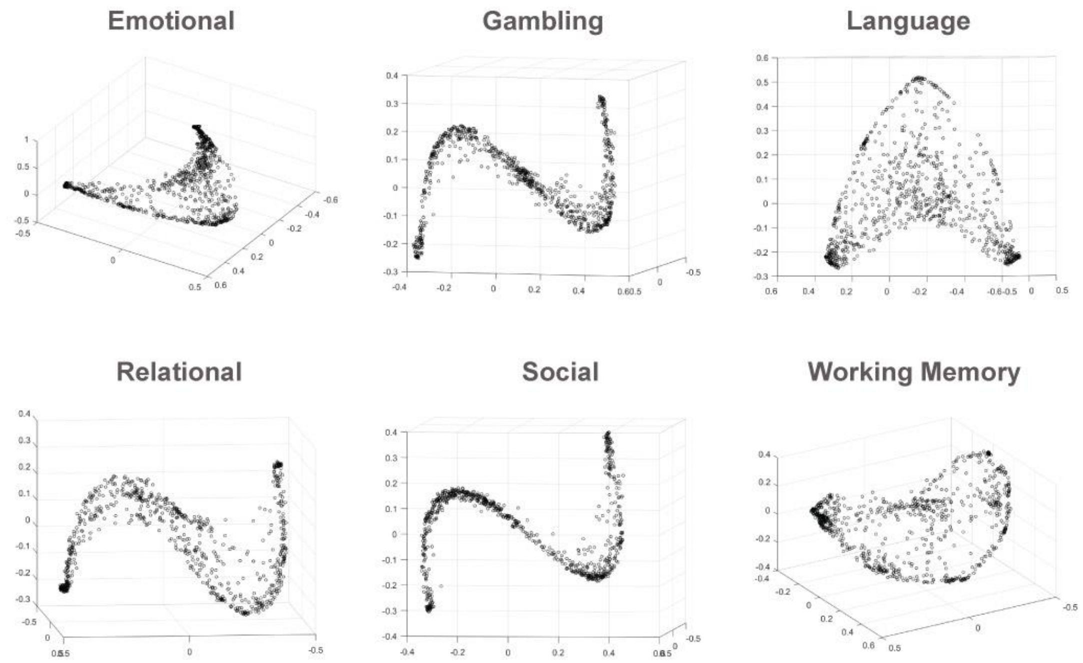


**Supplemental Figure 14**: Participants mapped into 3D space based on their scores derived from non-metric multidimensional scaling (MDS) of the clustering probability matrix. In order to allow for greater complexity of the data, 10 dimensions were calculated, but the first three are plotted. MDS produced components which were very similar to the PCA and recreated the observed ‘snake’ and ‘tortilla’ shapes.

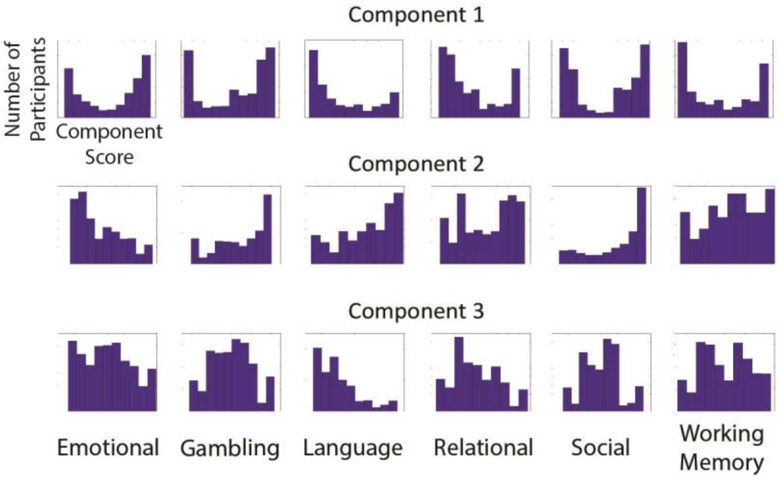


**Supplemental Figure 15**: Histograms of the top three components from the cluster bootstrap matrices for k=4.

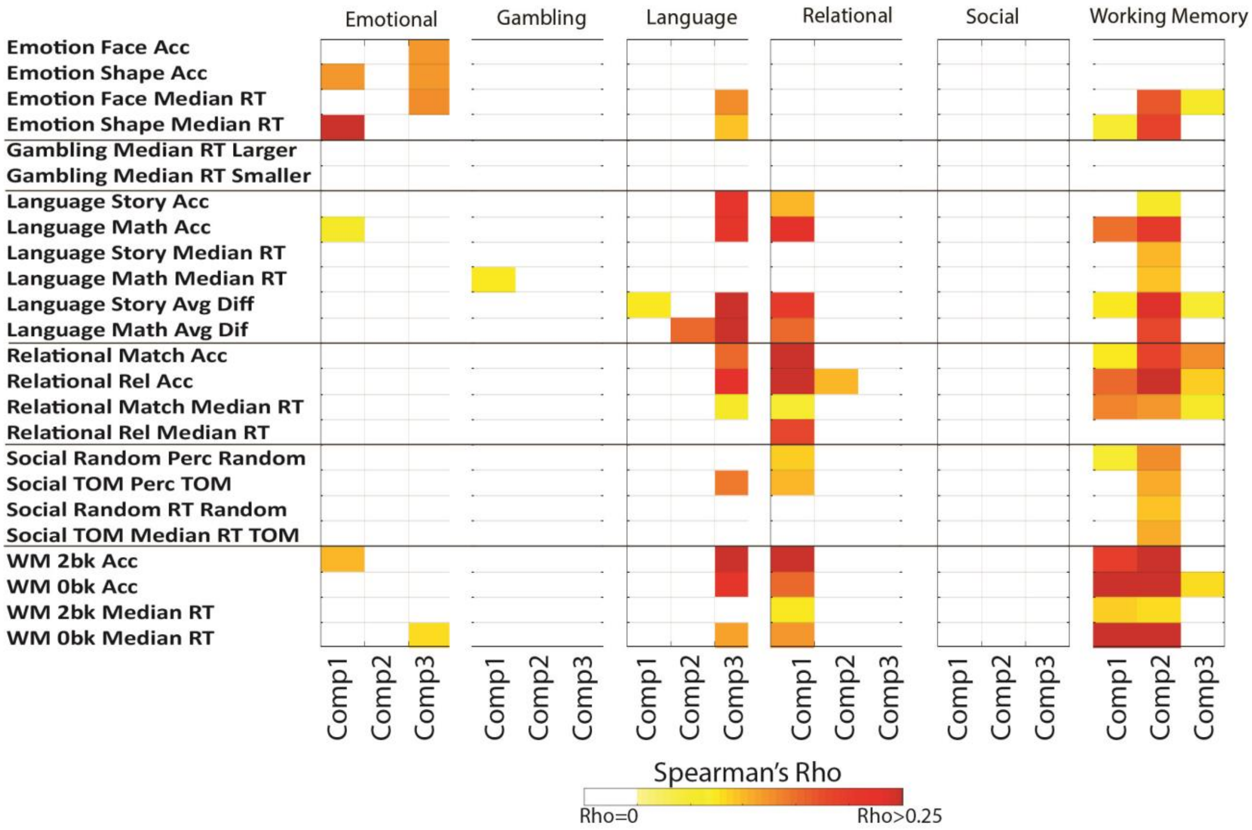


**Supplemental Figure 16**: Spearman's correlations between the first three principal components of the clustering probability matrices and performance on the fMRI tasks. For each fMRI task (large boxes) a Spearman's correlation was calculated for each component (columns with boxes, labeled as Comp1, Comp2, and Comp3, respectively) extracted from the clustering probability matrix for that task at k=4. Only correlations surviving FDR correction are shown in color (non-significant correlations are white). Because the directionality (positive-negative) of the component score is arbitrary, all correlations are presented as positive.
